# On the flexibility of basic risk attitudes in monkeys

**DOI:** 10.1101/282566

**Authors:** Shiva Farashahi, Habiba Azab, Benjamin Hayden, Alireza Soltani

## Abstract

Monkeys and other animals appear to share with humans two risk attitudes predicted by prospect theory: an inverse-S-shaped probability weighting function and a steeper utility curve for losses than for gains. These findings suggest that such preferences are stable traits with common neural substrates. We hypothesized instead that animals tailor their preferences to subtle changes in task contexts, making risk attitudes flexible. Previous studies used a limited number of outcomes, trial types, and contexts. To gain a broader perspective, we examined two large datasets of male macaques’ risky choices: one from a task with real (juice) gains and another from a token task with gains and losses. In contrast to previous findings, monkeys were risk-seeking for both gains and losses (i.e. lacked a reflection effect) and showed steeper gain than loss curves (loss-seeking). Utility curves for gains were substantially different in the two tasks. Monkeys showed nearly linear probability weightings in one task and S-shaped ones in the other; neither task produced a consistent inverse-S-shaped curve. To account for these observations, we developed and tested various computational models of the processes involved in the construction of reward value. We found that adaptive differential weighting of prospective gamble outcomes could partially account for the observed differences in the utility functions across the two experiments and thus, provide a plausible mechanism underlying flexible risk attitudes. Together, our results support the idea that risky choices are flexibly constructed at the time of elicitation and place important constraints on neural models of economic choice.

## INTRODUCTION

Humans and other animals live in a complex world in which uncertainty is often unavoidable (Kacelnik and Bateson, 1997; Pearson et al., 2014; Platt and Huettel, 2008). Understanding the strategies used to deal with risk –which we call risk attitudes – as well as underlying neural mechanisms is an important quest for behavioral economics, comparative psychology, foraging theory, and neuroscience (Kahneman and Tversky, 2000; McCoy and Platt, 2005; O’Neil and Schultz, 2010; Paglieri et al., 2014; So and Stuphorn, 2010; Trepel et al., 2005). When a strategy for dealing with risk is beneficial, it is liable to become selected for and canalized; i.e., become robustly seen across contexts and developmental trajectories. Consistent preferences across many or all members of a species have been often used to suggest that those preferences are innate and rely on similar neural substrates (e.g., Heilbronner et al., 2008; Heilbronner, 2017; Mendelson et al., 2016; De Petrillo et al., 2015; Stevens et al., 2005).

The rhesus macaque is a particularly important model organism in neuroeconomics. Macaques share many economic biases and preferences with humans, including attitudes towards counterfactual outcomes, the hot-hand effect, a peak-end bias, framing, cognitive dissonance, and the experience-description gap (Abe and Lee, 2011; Beran et al., 2014; Blanchard et al., 2014; Blanchard et al., 2015; Egan et al., 2007; Hayden et al, 2009; Heilbronner and Hayden, 2016; Lakshminarayanan et al., 2011). Some recent research suggests that macaques and other non-human primates share core risk attitudes as characterized by prospect theory (Kahneman and Tversky, 1979). Most notably, these include loss aversion (overweighting of possible losses compared to gains, Chen et al., 2006), the reflection effect (simultaneous risk-aversion with gains and risk-seeking with losses, Lakshmirayanan, et al., 2011), and an inverse-S-shaped probability weighting function (overweighting and underweighting of small and large probabilities, respectively; Stauffer et al., 2015). However, whereas humans are reliably risk-averse in many contexts, macaques are generally risk-seeking (Heilbronner and Hayden, 2013; but see Yamada et al., 2013). The consistency of these results across studies and, with the exception of risk-seeking, across species, have been used to suggest that such preferences are stable traits with common neural substrates and to motivate the use of non-human primates for studying choice under risk and uncertainty (Heilbronner, 2017).

While risk attitudes are important, cognitive flexibility is important for any organism that will encounter dynamic environments (Diamond, 2013). Flexible cognition that allows for rapid adjustment of risky choice strategy to even subtle changes in the environment should be selected for as well. Cognitive flexibility is not necessarily inconsistent with evolved risk attitudes, but primates’ remarkable flexibility raises the possibility that ostensibly shared risk attitudes may be task-dependent. Specifically, if preferences are task-dependent, then comparing two species’ attitudes in the same task, or one species’ attitudes across two tasks may produce similar preferences because the computational demands of the task or tasks are similar (e.g., range of reward probabilities). For this reason, testing risk attitudes across multiple contexts can be informative.

To obtain a broader view on flexibility of risk attitudes, we examined two large datasets supplemented with new data: one from a juice-based gambling task in which monkeys chose between two win/nothing gambles on each trial (Strait et al., 2014; Strait et al., 2015), and one from a token-based gambling task in which monkeys selected between two mixed (win/loss) gambles (Azab and Hayden, 2017, 2018; Strait et al., 2016). Our aim was to examine monkeys’ behavior in light of extant findings and predictions made under prospect theory. By fitting choice behavior with various models using cross-validation, we found that monkeys were risk-seeking in both tasks, although their utility curves for gains had different convex shapes. Monkeys were loss-seeking in the token-gambling task and exhibited a convex utility curve for losses shallower than the one for gains. Finally, the probability weighting function was S-shape (the inverse of the previously reported shape) in the juice-gambling task and almost linear in the token-gambling task.

## MATERIALS AND METHODS

### Overview of the experimental procedures

Behavioral data were collected in two separate experiments in which monkeys selected between two gambles offering juice or tokens. In each trial of the juice-gambling experiment, monkeys selected one of two options, each offering a simple gamble for juice or water (Strait et al., 2014; Strait et al., 2015). Options were represented by a rectangular bar and offered either a gamble or a safe bet (100% probability) for liquid reward. Gamble offers were represented by a bar that was divided into two portions corresponding to the two possible outcomes: no reward and a medium or large reward (**Figure 1a**).

**Figure 1.**
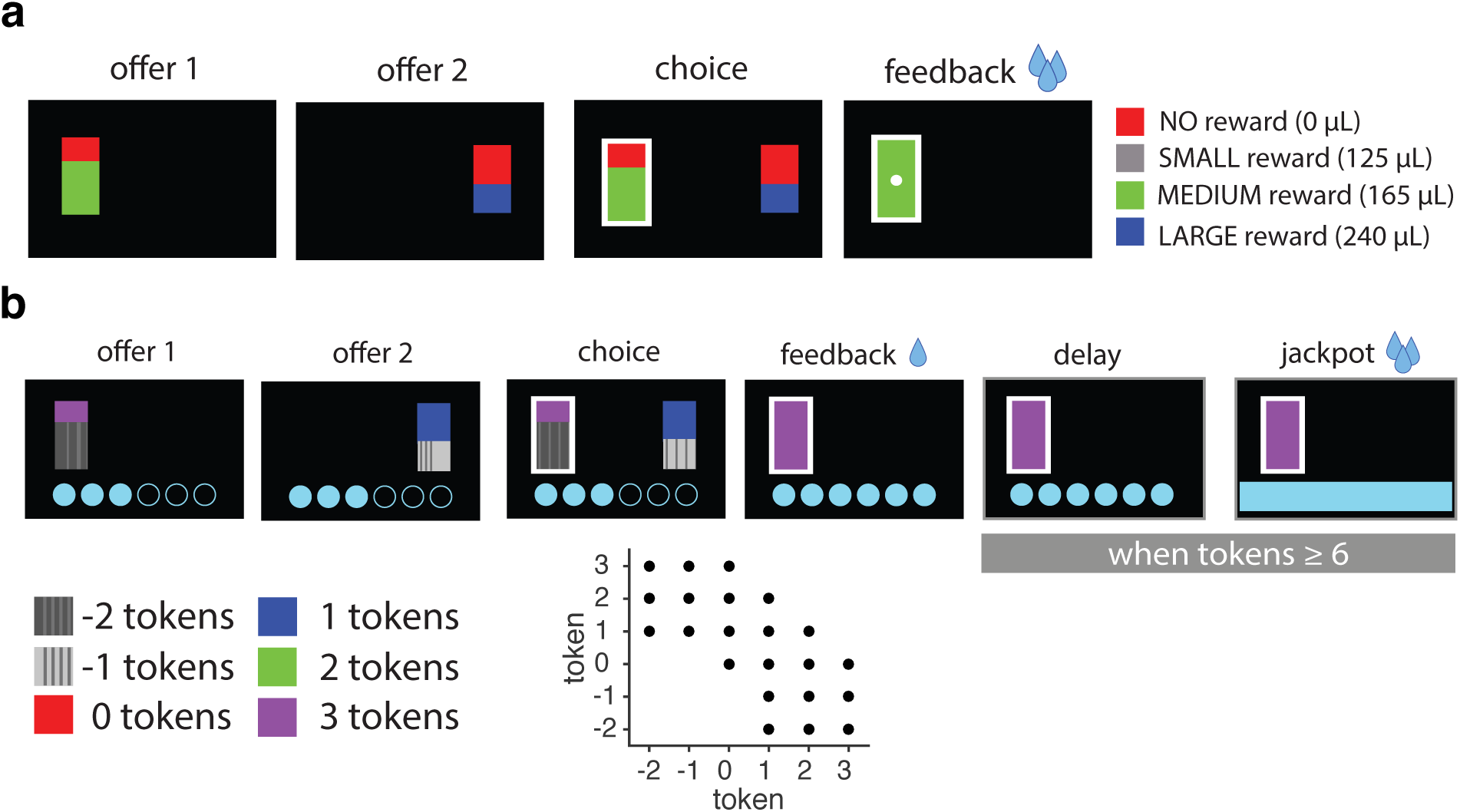
Experimental procedure. (**a**) Timeline of the juice-gambling task. In each trial, two options were presented, each offering a gamble for juice reward. Gambles were represented by a rectangle, some portion of which was red, blue, or green, signifying no reward, medium, or large reward, respectively. The area of the colored portion indicates the probability that choosing that offer would yield the corresponding reward. We also used a safe offer that was entirely gray and always carried a 100% probability of a small reward. (**b**) Timeline of the token-gambling task with gains and losses. In each trial, two options were presented, each offering a gamble for tokens. The size of each colored portion within each offer indicated the probability that choosing that offer would yield the corresponding outcome. A small reward was administered for each completed trial. When at least six tokens were earned, a large “jackpot” reward was administered and the earned token count was reset to 0. The inset shows the colors associated with different tokens and combinations used.

In each trial of the token-gambling experiment, monkeys selected between two options, each offering a mixed-gamble for tokens (Strait et al. 2016; Azab and Hayden, 2017, 2018). Visual display of gambles was similar to the juice-gambling task except six colors were used corresponding to six possible reward magnitudes in terms of tokens (three gains, two losses, and zero; **Figure 1b**). In addition, the probabilities of reward outcome were limited to five values (0.1, 0.3, 0.5, 0.7, 0.9). Each gamble included at least one positive or zero-outcome, ensuring that every gamble carried the possibility of a win. This decreased the number of trivial choices presented to subjects, and maintained motivation. Monkeys were trained to collect six tokens to receive a large (300*μ*L) liquid reward (see *token-gambling task* below for more details). Therefore, each token corresponded to 50*μ*L of reward juice.

In total, three male monkeys (subject B, C, and J) performed 108,272 and 66,500 trials in the juice and token-gambling tasks, respectively. Monkeys B and J participated in both experiments. Monkeys B, C, and J performed 70,700, 24,700, and 12,872 trials in the juice-gambling task, respectively. Monkeys B and J performed 28,700 and 37,800 trials in the token-gambling task, respectively. Subjects were initially trained on a two-option task (Strait et al., 2014) and then later also trained with a task that involved single-option accept-reject gambles (Blanchard et al., 2015b). Although subjects were not tested with novel colors in this study, we have extensively tested macaques’ abilities to learn new associations quickly. This approach to training risk tasks was explained in detail elsewhere (Hayden, Heilbronner, and Platt, 2010).

Proportional gambling tasks have been used by many labs since 2010 (e.g. O’Neill and Schultz, 2010; So and Stuphorn 2012; Yamada and Glimcher, 2013; Strait and Hayden, 2014; Chen and Stuphorn, 2015). There is plentiful evidence that monkeys readily understand and correctly interpret such displays with no special training requirements. The Hayden lab has been developing methods for training macaques to perform such tasks for over a decade and we have developed several checks and training strategies to make sure they understand the task. Subjects were trained in two stages. Our subjects were first trained extensively (for two or more years) on a simple gambling task with multiple possible juice (i.e. non-token) reward amounts. In this stage, they were tested on multiple variations of the gambling task, and performance was validated through multiple control tests (Hayden and Heilbronner, 2010). Performance was consistent (including two consistent biases, risk-seeking and win-stay-lose-shift) across single option (Blanchard et al., 2015b) and two-option (Strait et al., 2014) versions of the task. The token element of the task was new to our lab, although it has been used in other labs before (e.g. Seo and Lee, 2009; Seo et al., 2014). Behavior in the token version of the task was overall quite similar to that in the juice version, indicating that the monkeys readily learned to treat secondary rewards as reinforcing. However, the strongest evidence for the monkeys’ understanding of the task comes from their consistent preferences for higher probabilities of large rewards and smaller probabilities of small rewards.

### Juice-gambling task

Two offers were presented on each trial. Each offer was represented by a rectangle 300 pixels tall and 80 pixels wide (11.35° of visual angle tall and 4.08° of visual angle wide). Options offered either a gamble or a safe (100% probability) bet for liquid reward. Gamble offers were defined by two parameters, reward size and probability. Each gamble rectangle was divided into two portions: one red and the other either blue or green. The size of the blue or green portions signified the probability of winning a medium (mean 165 *μ*L) or large reward (mean 240 *μ*L), respectively. These probabilities were drawn from a uniform distribution between 0 and 100%. The rest of the bar was colored red; the size of the red portion indicated the probability of no reward. The safe offer was entirely gray, and always carried a 100% probability of a small reward (125 *μ*L).

On each trial, one offer appeared on the left side of the screen and the other appeared on the right. Offers were separated from the fixation point by 550 pixels (27.53° of visual angle). The side of the first and second offer (left and right) was randomized by trial. Each offer appeared for 400 ms and was followed by a 600 ms blank period. Monkeys were free to fixate upon the offers when they appeared (and in our casual observations almost always did so). After the offers were presented separately, a central fixation spot appeared and the monkey fixated on it for 100 ms. Following this, both offers appeared simultaneously and the animal indicated its choice by shifting gaze to its preferred offer and maintaining fixation on it for 200 ms. Failure to maintain gaze for 200 ms did not lead to the end of the trial, but instead returned the monkey to a choice state; thus monkeys were free to change their mind if they did so within 200 ms (although in our observations, they seldom did so). Following a successful 200-ms fixation, the gamble was immediately resolved and reward delivered. Trials that took more than 7 seconds were considered inattentive trials and were not included in analysis (this removed <1% of trials). Outcomes that yielded rewards were accompanied by a visual cue: a white circle in the center of the chosen offer. All trials were followed by an 800-ms inter-trial interval with a blank screen.

### Token-gambling task

Monkeys performed a mixed (two-option) gambling task. The task was similar to one we have used previously (Strait et al. 2014; Strait et al. 2015), albeit with two major differences: first, monkeys gambled for virtual tokens —rather than liquid —rewards, and, second, outcomes could be losses as well as wins.

Two offers were presented on each trial. Each offer was represented by a rectangle 300 pixels tall and 80 pixels wide (11.35° of visual angle tall and 4.08° of visual angle wide). 20% of options were safe (100% probability of either 0 or 1 token), while the remaining 80% were gambles. Safe offers were entirely red (0 tokens) or blue (1 token). The size of each portion indicated the probability of the respective reward. Each gamble rectangle was divided horizontally into a top and bottom portion, each colored according to the token reward offered. Gamble offers were thus defined by three parameters: two possible token outcomes, and probability of the top outcome (the probability of the bottom was strictly determined by the probability of the top). The probability of the outcome was selected from the following values: 0.1, 0.3, 0.5, 0.7, 0.9. The token values of the two possible outcomes were selected at random from the values −2 (black stripe), −1 (gray stripe), 0 (red), 1 (blue), 2 (green), or 3 (purple). The combinations used are shown in the inset of **Figure 1b**. Only red (0 token) and blue (1 token) were used as safe offers.

Six initially unfilled circles arranged horizontally at the bottom of the screen indicated the number of tokens to be collected before the subject obtained a liquid reward. These circles were filled appropriately at the end of each trial, according to the outcome of that trial. When 6 or more tokens were collected, the tokens were covered with a solid rectangle while a liquid reward was delivered. Tokens beyond 6 did not carry over, nor could number of tokens fall below zero.

On each trial, one offer appeared on the left side of the screen and the other appeared on the right. Offers were separated from the fixation point by 550 pixels (27.53° of visual angle). The side of the first offer (left and right) was randomized by trial. Each offer appeared for 600 ms and was followed by a 150 ms blank period. Monkeys were free to fixate upon the offers when they appeared (and in our observations almost always did so). After the offers were presented separately, a central fixation spot appeared and the monkey fixated on it for 100 ms. Following this, both offers appeared simultaneously, and the animal indicated its choice by shifting gaze to its preferred offer and maintaining fixation for 200 ms. Failure to maintain gaze for 200 ms did not lead to the end of the trial, but instead returned the monkey to a choice state; thus, monkeys were free to change their mind if they did so within 200 ms (although in our observations, they seldom did so). A successful 200 ms fixation was followed by a 750 ms delay, after which the gamble was resolved and a small ‘motivation’ reward (100 *μ*L) was delivered, regardless of the outcome of the gamble, to sustain motivation. This small reward was delivered within a 300 ms window. If 6 tokens were collected, a delay of 500 ms was followed by a large liquid reward (300 *μ*L) within a 300 ms window, followed by a random inter-trial interval (ITI) between 500 and 1500 ms. If 6 tokens were not collected, subjects proceeded immediately to the ITI.

Each gamble included at least one positive or zero-outcome, ensuring that every gamble carried the possibility of a win. This decreased the number of trivial choices presented to subjects and maintained motivation.

### Surgical procedures, eye tracking and reward delivery

All procedures were approved by the University Committee on Animal Resources at the University of Rochester and were designed and conducted in compliance with the Public Health Service’s Guide for the Care and Use of Animals. Three male rhesus macaques (Macaca mulatta) served as subjects. A small prosthesis for holding the head was used. A Cilux recording chamber (Crist Instruments) was placed over the prefrontal cortex. Animals were habituated to laboratory conditions and then trained to perform oculomotor tasks for liquid reward. Animals received appropriate analgesics and antibiotics after all procedures. Throughout both behavioral and physiological recording sessions, the chamber was kept sterile with regular antibiotic washes and sealed with sterile caps. All recordings were performed during the animals’ light cycle between 8 a.m. and 5 p.m.

Eye position was sampled at 1,000 Hz by an infrared eye-monitoring camera system (SR Research). Stimuli were controlled by a computer running Matlab (Mathworks) with Psychtoolbox (Brainard 1997) and Eyelink Toolbox (Cornelissen et al. 2002). Visual stimuli were colored rectangles on a computer monitor placed 57 cm from the animal and centered on its eyes (**Figure 1a-b**). A standard solenoid valve controlled the duration of juice delivery. The relationship between solenoid open time and juice volume was established and confirmed before, during, and after recording.

### Overview of computational models

We first used four base models (EV, EV+PW, EU, and SU) for the estimation of subjective value. In all these models, the subjective value of each gamble (say the gamble on the left) was computed as follows:

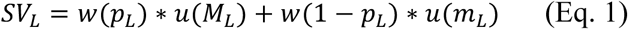

where *SV_L_* is the subjective value of the left gamble, *M_L_* and *p_L_* are the magnitude (in *μ*L) and probability associated with the left gamble’s larger magnitude outcome, *m_L_* is the magnitude of the other left gamble outcome (*M_L_* > *m_L_*) which is equal to zero in the juice-gambling task, *u*(*m*) is the utility function (UF), and *w*(*p*) is the probability weighting function (PWF). The four models differed in the form of their utility and probability weighting functions. The EV model included linear utility and probability weighting functions. The EU model included only a nonlinear utility function whereas the EV+PW included only a nonlinear probability weighting function. Finally, the SU included both nonlinear utility and probability weighting functions (see *base models* below for more details).

The estimated subjective values of the two options presented in each trial were then used to compute the probability of selecting between the two options based on a logistic function

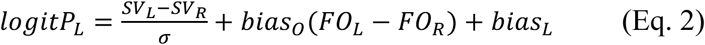

where *P_L_* is the probability of choosing the left option, *bias_L_* measures a response bias toward the left option to capture any location bias, *bias_o_* measures a response bias toward the first offer that appeared on the screen (order bias) and was only significant in the token-gambling task (*FO_L_*(*FO_R_*) is 1 if the first offer appeared on the left (right) side), and *σ* is a parameter that measures the level of stochasticity in decision processes.

We also extended our base models to include two types of differential weighting mechanisms (see *Models with differential weighting mechanism* below for more details). First, we considered alternative ‘within-option’ differential weighting mechanisms by which the gamble outcome with a larger reward magnitude, reward probability, or expected value could influence the overall value more than the alternative outcome. This was done to investigate how magnitudes and probabilities of the two possible gamble outcomes can influence the weight of each gamble outcome on the overall gamble value. These models were only used for the token-gambling task since gambles in the juice-gambling task only had one non-zero outcome. Second, we considered the possibility that when comparing two gambles, the value of the better outcome of each gamble (in terms of magnitude, probability, or EV) could influence their overall value relative to the other gamble (‘cross-option’ differential weighting). This was done to investigate how non-zero (or the better) outcomes of the gambles on each trial modulate the value of these gamble in the juice- (respectively, token-) gambling task.

### Base models

In the expected value (EV) model, actual probabilities and a linear utility function were used to estimate the subjective value of each gamble. However, this model also includes different slopes for gains and losses as follows:

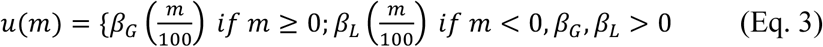

where *β_G_* and *β_L_* are slopes for the gain and loss domains, respectively. We normalized the juice reward magnitude by 100*μ*L in order to limit utility to small numbers.

In the expected utility (EU) model we considered a nonlinear utility function (UF) and a loss-aversion factor as follows:

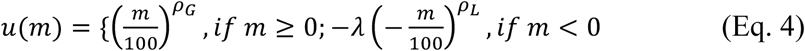

where *u*(*m*) is the subjective utility, *λ* is the loss-aversion factor, and *ρ_G_* and *ρ_L_* are the exponents of the power law function and determine risk-aversion for the gain and loss domains, respectively; *ρ* > 1 indicates risk-seeking, *ρ* < 1 indicates risk-aversion, and *ρ* = 1 indicates risk-neutrality.

In the EV+PW model, we considered a linear utility function and nonlinear probability weighting function (PWF). The PWF was computed using a 1-parameter Prelec function as follows:

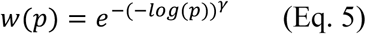

where *w*(*p*) is the PWF, and *γ* is a parameter that determines probability distortion. In general, *γ* > 1 indicates risk-aversion, *γ* < 1 indicates risk-seeking, and *γ* = 1 indicates risk-neutrality. Finally, in the SU model, we used both nonlinear utility and nonlinear probability weighting functions to estimate the subjective value of each gamble.

### Models with differential weighting mechanisms

We extended our base models to include two types of differential weighting mechanisms. First, we considered alternative differential weighting mechanisms by which the gamble outcome with a larger reward magnitude, reward probability, or expected value could influence the overall value more than the alternative outcome (‘within-option’ differential weighting; **Figure 2a-c**). Second, we considered the possibility that when comparing two gambles, the value of the better outcome of each gamble (in terms of magnitude, probability, or EV) could influence their overall value relative to the other gamble (‘cross-option’ differential weighting; **Figure 2d-i**).

**Figure 2.**
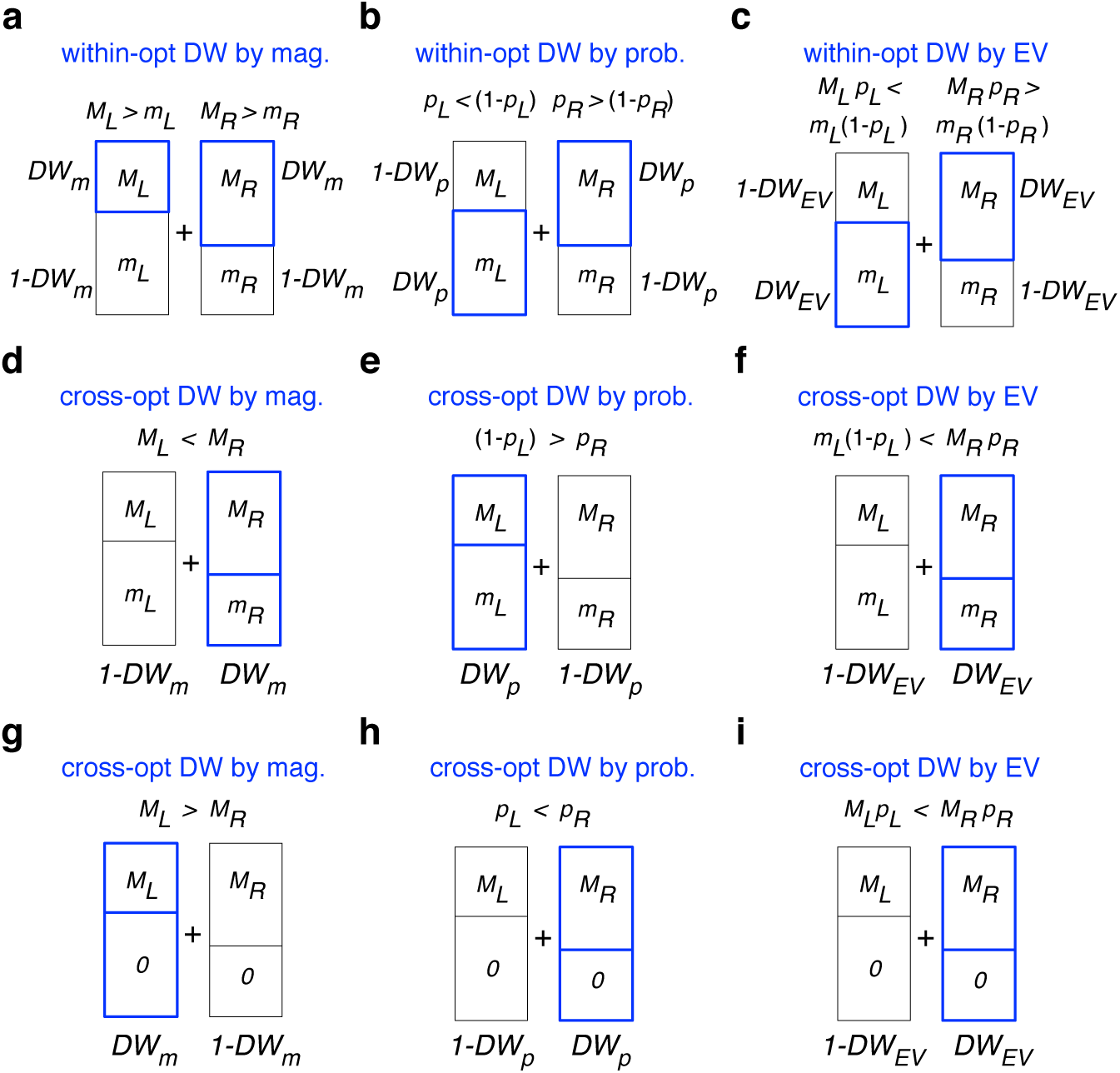
Alternative models for differential weighting. (**a-c**) Alternative differential weighting between the two outcomes of each gamble (within-option) in the token-gambling task with two non-zero reward outcomes. Panels a-c show three mechanisms for how magnitudes and probabilities of the two possible gamble outcomes can influence the weight of each gamble outcome on the overall gamble value: differential weighting (DW) by magnitude (a); differential weighting by probability (b); and differential weighting by EV (c). In all panels, *M_L_* and *p_L_* indicate the magnitude and probability associated with the left gamble’s larger magnitude outcome, respectively, and *m_L_* is the magnitude of the other left gamble outcome (the probability of this outcome is 1 − *p_L_*). The same convention is used for the right gamble. The blue box shows the outcome that is assigned with a larger weight based on a given mechanism. The DW factors determine the strength of differential weighting according to the reward magnitude (*DW_m_*), reward probability (*DW_p_*), and expected value (*DW_EV_*) of the two outcomes. (**d-f**) Alternative differential weighting between better outcomes of the two alternative gambles (cross-option) in the token-gambling task with two non-zero reward outcomes. Panels d-f show three mechanism for how magnitudes and probabilities of the better outcome of the two alternative gambles can modulate their values: cross-option differential weighting by magnitude (d), cross-option differential weighting by probability (e), and cross-option differential weighting by EV (f). The blue box shows the gamble that is assigned with a larger weight based on a given mechanism. The DW factors determine the strength of differential weighting according to (d) the reward magnitude, (e) reward probability, and (f) expected value of the two gambles. (**g-i**) Alternative cross-option differential weighting between non-zero outcomes of the two alternative gambles in the juice-gambling task. Panels (g-i) show three mechanisms for how magnitudes and probabilities of the non-zero outcome of the two alternative gambles can modulate their values. Importantly, as shown in **Figures 2-1** and **2-2**, our fitting method is able to correctly identify the model used to generate a given set of data and thus can distinguish between the alternative models.

In all models with within-option differential weighting, the subjective value of a gamble was computed as follows:

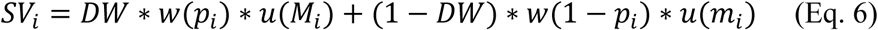

where *DW*(differential-weighting factor) determines the strength of differential weighting by one of the three alternative mechanisms: differential weighting by magnitude; differential weighting by probability; and differential weighting by EV (**Figure 2a-c**).

We constructed three within-option differential weighting models (differential weighting by magnitude, differential weighting by probability, and differential weighting by EV) to investigate how magnitudes and probabilities of the two possible gamble outcomes can influence the weight of each gamble outcome on the overall gamble value. These models were only used for the token-gambling task since gambles in the juice-gambling task only had one non-zero outcome.

In the model with within-option differential weighting by magnitude (diff. weight by mag.), the subjective value of each gamble (say for the left gamble) was computed as follows:

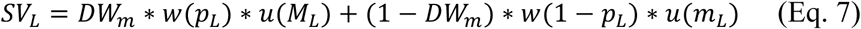

where *DW_m_* (differential-weighting factor) determines the strength of differential weighting by magnitudes, and *M_L_* and *p_L_* are the magnitude and probability associated with the left gamble’s larger magnitude outcome, respectively, and *m_m_* is the magnitude of the other left gamble outcome (*M_L_* > *m_L_*).

In the model with within-option differential weighting by probability (diff. weight by prob.), the subjective value was computed as follows:

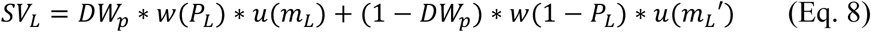

where *DW_p_* determines the strength of differential weighting by probability, *m_L_* (*m_L_*′) is the magnitude associated with the left gamble’s larger (respectively, smaller) probability outcome (*P_L_* > 0.5).

Finally, in the model with within-option differential weighting by expected value (diff. weight by EV), the subjective value was computed as follows:

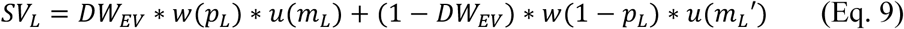

where *DW_EV_* determines the strength of differential weighting by expected value, *m_L_* and *p_L_* are the magnitude and probability associated with the left gamble’s outcome with a larger expected value, respectively, and *m_L_*′ is the magnitude associated with the left gamble’s outcome with a smaller expected value (*p_L_* ∗ *m_L_* > (1 − *p_L_*) ∗ *m_L_*′).

We also constructed three cross-option differential weighting models (cross-opt differential weighting by magnitude, cross-opt differential weighting by probability, and cross-opt differential weighting by EV) to investigate how non-zero (or the better) outcomes of the gambles on each trial modulate the value of these gamble in the juice-gambling (respectively, token-gambling) task.

In all the models with cross-option differential weighting, the probability of selecting between the gambles was computed as follows (if the left gamble was assigned with the larger weight):

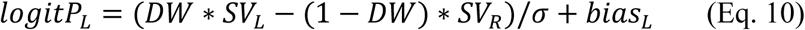

where *DW* determines the strength of differential weighting between the two non-zero or better outcomes of the two alternative gambles based on one of the three alternative mechanisms: differential weighting by magnitude; differential weighting by probability; and differential weighting by EV (**Figure 2d-i**).

In the models with cross-option differential weighting by magnitude (**Figure 2g**), *DW* was multiplied by the value of the gamble with a larger magnitude outcome; for example the left gamble, if *M_L_* > *M_R_*, where *M_i_* denotes the larger magnitude outcome of each gamble. In the models with cross-option differential weighting by probability (**Figure 2h**), *DW* was multiplied by the value of the gamble with a larger probability outcome for which the magnitude was non-zero; for example the left gamble, if *P_L_* > *P_R_*, where *P_i_* denotes the larger probability outcome of each gamble. Finally, in the models with cross-option differential weighting by expected value (**Figure 2i**), *DW* was multiplied by the value of the gamble with a larger expected value outcome; for example the left gamble, if *m_L_* ∗ (1 − *p_L_*) > *M_R_* ∗ *p_R_*, when *m_L_* ∗ (1 − *p_L_*) and *M_R_* ∗ *p_R_* are the EV of the larger EV outcomes of the two gambles.

### Fitting procedure and data analyses

To examine how monkeys constructed subjective value for risky options, we used various models to fit choice behavior during each gambling task; the best model revealed the most plausible mechanism for the construction of subjective value for a given monkey/task. Models were fitted to experimental data by minimizing the negative log likelihood of the predicted choice probability given different model parameters using the fminsearch function in MATLAB (Mathworks). There are two main issues when comparing the goodness-of-fit between models with different number of parameters: more complex models could explain data better by virtue of having a greater number of parameters; models with more parameters could over-fit the data such that the fitting is not generalizable to similar datasets. For these reasons, we fit choice behavior with different models based on a 5-fold cross-validation method, using parameters estimated from 80% of the data for a given monkey/task to predict choices on the remaining 20%. Importantly, cross-validation automatically deals with different numbers of model parameters because redundant parameters result in over-fitting and thus do not add any explanatory power. Moreover, it has been shown that in many cases, the cross-validation method provides an approximation to the Akaike information criterion (AIC) whereas the AIC does not address the over-fitting issue. The cross-validation was done 50 times separately for data from each monkey in a given task.

In addition, we also fit choice behavior from each session of the experiment individually in order to capture the diversity of risk attitudes on different days of the experiment. We used interquartile range rule to remove outlier sessions in terms of the estimated parameters. More specifically, we only included sessions that did not yield an outlier for any of the fitting parameters. This was done to ensure a reliable estimate for all the parameters in a given session. In the juice-gambling task, the exclusion criterion resulted in removal of 2% and 11% of sessions from the lower and upper outlier bounds, respectively. In the token-gambling task, this exclusion criterion resulted in removal of 4% and 12% of sessions from the lower and upper outlier bounds, respectively. Importantly, we obtained qualitatively similar results for session-by-session analyses even with the inclusion of outlier sessions.

To test whether our fitting procedure is able to distinguish between alternative models and identify the correct model and to accurately estimate model parameters, we simulated the aforementioned sixteen models over a range of parameters estimated from monkeys’ choice behavior in the two experiments. More specifically, we generated choice data for the juice-gambling task using the exponent of the utility function (*ρ*) ranging from 1 to 4, the probability distortion parameter (*γ*) ranging from 0.8 to 2, the differential-weighting factor (*DW*) ranging from 0.55 to 0.65 and the stochasticity in choice (*σ*) ranging from 0.5 to 10. In the token-gambling task, we generated choice data by adopting the following range for model parameters: [1, 2] for the exponent of the utility function between (*ρ*), [0.4, 1.2] for the loss aversion factor (*λ*), [0.8, 1.2] for the probability distortion parameter (*γ*), [0.55, 0.65] for the differential weighting (*DW*), and [0.4, 1] for stochasticity in choice (*σ*). We then fit the simulated data with all the models to compute the goodness-of-fit (in terms of AIC) and to estimate model parameters. Because model parameters could take on very different values, we computed the error in estimation of model parameters using the relative value of each estimated parameter to its actual value. The average goodness-of-fit and estimation error were calculated by averaging the corresponding values over all fits based on all sets of parameters. Moreover, in order to account for the overall difficulty of fitting data generated with certain models, we rescaled AIC values across all models used to fit a given set of simulated data. This rescaling was done by first subtracting the minimum AIC value obtained by fitting a given set of data and then dividing the outcome by the difference between the maximum and minimum values of AIC for that set of data.

To test correlation between model parameters, we used two methods: the Hessian matrix and session-by-session values of fitting parameters. First, we numerically estimated correlations between model parameters using the Hessian matrix for the SU model and the SU model with differential weighting at parameter values for which we obtained of best fit. The relationship between the matrix of correlation between model parameters and the Hessian matrix builds on the theorem that because the maximum likelihood estimator is asymptotically normal, the distribution of the maximum likelihood estimator (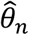) can be approximated by a multivariate normal distribution with a certain mean (*θ*_0_) and a covariance matrix 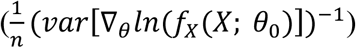, where *n* is the number of model parameters. This covariance matrix can be estimated by 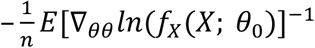, where 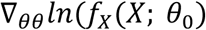 is the matrix of the second-order partial derivatives of the log-likelihood function, or the Hessian matrix. As a result, the matrix of correlation between model parameters can be calculated from the inverse of the Hessian matrix. To estimate the Hessian matrix, we first computed the derivatives of the log likelihood with respect to model parameters to form the Jacobian matrix. Next, we calculated the derivative of the Jacobian matrix to compute the Hessian matrix of the cost function for fitting.

Second, we directly calculated the correlation between model parameters based on the estimated parameters across all sessions using the base SU model and the SU model with differential weighting. Using session-by-session fitting parameters, we also calculated the correlation between model parameters of both models. These two methods for calculating correlation between model parameters yield compatible results (see Results).

We also examined the likelihood surface of the model in order to calculate the error associated with the estimated parameters. We calculated variability in the estimate of negative log-likelihood function (by computing the standard deviation of this function, *std*(−*LL*)) at the global minimum across many instances of cross-validation. We then calculated the order of magnitude (scale) of the error associated with estimated parameters using the eigenvector associated with the smallest eigenvalue of the Hessian matrix (as a measure of the direction with the minimum slope of log-likelihood surface) as follows:

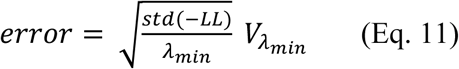

where *λ_min_* is the smallest eigenvalue and *V*_λ_*_min_* is the corresponding eigenvector.

Finally, to quantify changes in the sensitivity to reward information as a function of the number of collected tokens (**Figure 5**), we fit the psychometric function using a sigmoid function and estimates indifference point (*μ*) and stochasticity in choice (*σ*):

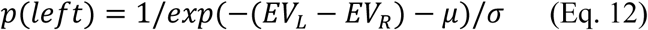

where *p*(*left*) is the probability of choosing the option on left, and *EV_L_* and *EV_R_* are the expected value of the left and right options, respectively.

## RESULTS

We used various computational models to analyze monkeys’ choice behavior from two separate experiments in which subjects chose between two options (gambles or safe options) offering either juice (juice-gambling task) or token (token-gambling task) rewards (**Figure 1**). The juice-gambling task involved options with the possibility of one of three reward sizes or no reward, whereas options in the token-gambling task involved a mix of gain, loss, or no reward possibilities (see Materials and Methods).

### Monkeys exhibit risk-seeking and loss-seeking

We first examined whether the animals appropriately integrated information about reward magnitude and probability to select between gambles. To do so, we computed the probability of choosing the left gamble as a function of the difference between the expected values (i.e. reward probability times magnitude) of the left and right gambles (**Figure 3c-d**). This analysis showed that all monkeys consistently selected the gamble with higher expected value (81%, 84% and 85% for monkeys B, C and J in the juice-gambling task and 79% and 74% for monkeys B and J in the token-gambling task; binomial test, *p* < 10^−5^). Moreover, psychometric functions plotted in **Figure 3c-d** provide strong evidence that all monkeys considered both length (probability) and color (magnitude) of gambles for making decisions.

**Figure 3.**
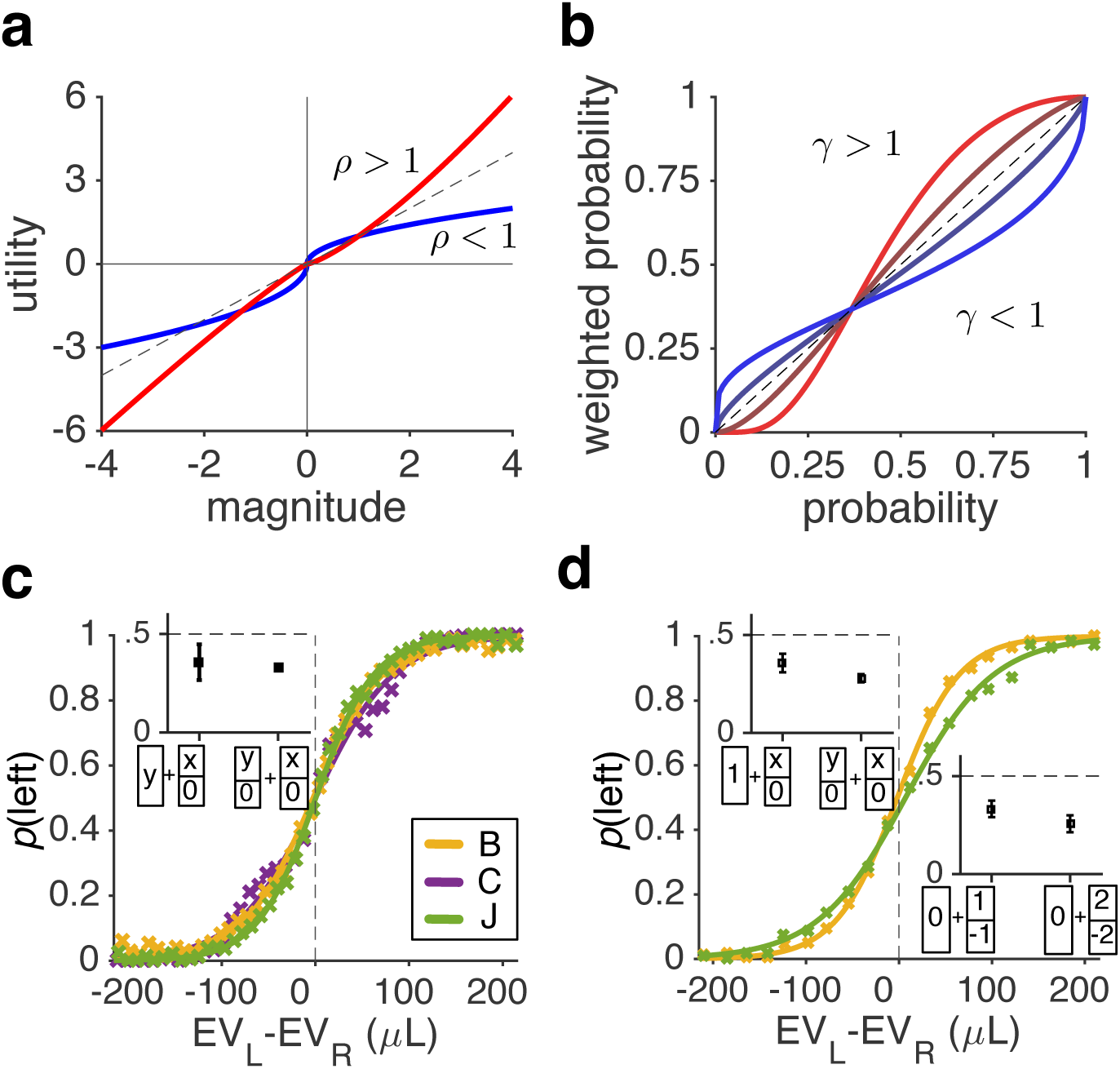
Standard and non-standard evaluation of reward magnitude and probability according to prospect theory, and the overall sensitivity of monkeys’ choice behavior to the difference in expected values of gambles on each trial. (**a**) Utility function quantifying the relationship between reward magnitude and subjective utility. Plotted in blue is a hypothetical, standard utility function based on prospect theory with concave and convex curves for gains and losses, respectively. In contrast, the red curve shows a utility function that is convex for gains and concave for losses, resulting in risk-seeking behavior for gains and risk-aversion for losses and thus respecting the reflection effect. Parameter *ρ* is the exponent of the power law used to generate the utility curves. (**b**) Probability weighting function quantifying the transformation of actual reward probability for making decisions. Plotted are several possible shapes of probability weighting. Prospect theory predicts inverse-S-shaped weighting functions (blue curves). Parameter *γ* determines the curvature of the function. (**c**) Psychometric functions in two monkeys during the juice-gambling task. Probability of choosing the left target is plotted as a function of the difference in expected values of two gambles in a given trial. The inset plots the probability (mean±s.e.m.) of choosing a sure option against a gamble with an equal expected value (*n* = 32 trials), and the probability of choosing the less risky option in pairs of gambles with equal expected values (*n* = 717 trials). (**d**) The same as in (c) but for the token-gambling task. The top inset plots the probability of choosing the sure option of one token against a gamble with an equal expected value (*n* = 69 trials), and the probability of choosing the less risky option in pairs of gambles with equal expected values (*n* = 271 trials). The bottom inset plots the probability of choosing the sure option with no reward over a 50/50 gamble of winning and losing one (*n* = 55 trials) or two tokens (*n* = 62 trials).

To measure overall preference for risk and loss, we next examined choices between pairs of a safe option and a risky gamble with equal expected value, or between pairs of risky gambles with equal expected values. We found that in both experiments, monkeys consistently selected the offer with the smaller reward probability and larger reward magnitude (i.e., the more risky option) over the one with the larger reward probability and smaller reward magnitude, indicating significant risk-seeking behavior (**Figure 3c-d insets**). Moreover, in the token-gambling task with gains and losses, monkeys consistently selected the 50/50 gambles with equal amounts of gains and losses over the sure option that did not deliver reward. This indicates that monkeys preferred to accept, rather than reject, gambles with loss and zero expected value, signifying loss-seeking behavior (**Figure 3d inset**).

To better demonstrate that monkeys understood the task and incorporated information about both reward probability and magnitude, we calculated choice probability separately for each set of gambles with similar reward magnitudes as a function of the probability of reward for the larger magnitude outcomes of the two gambles, or the only gamble when the competing choice option was a safe one (**Figure 4**). This analysis showed that the probability of choosing a gamble increased as the probability of reward for its larger magnitude outcome increased, indicating that monkeys did incorporate the length of a given colored portion (i.e. reward probability) in their choices.

**Figure 4.**
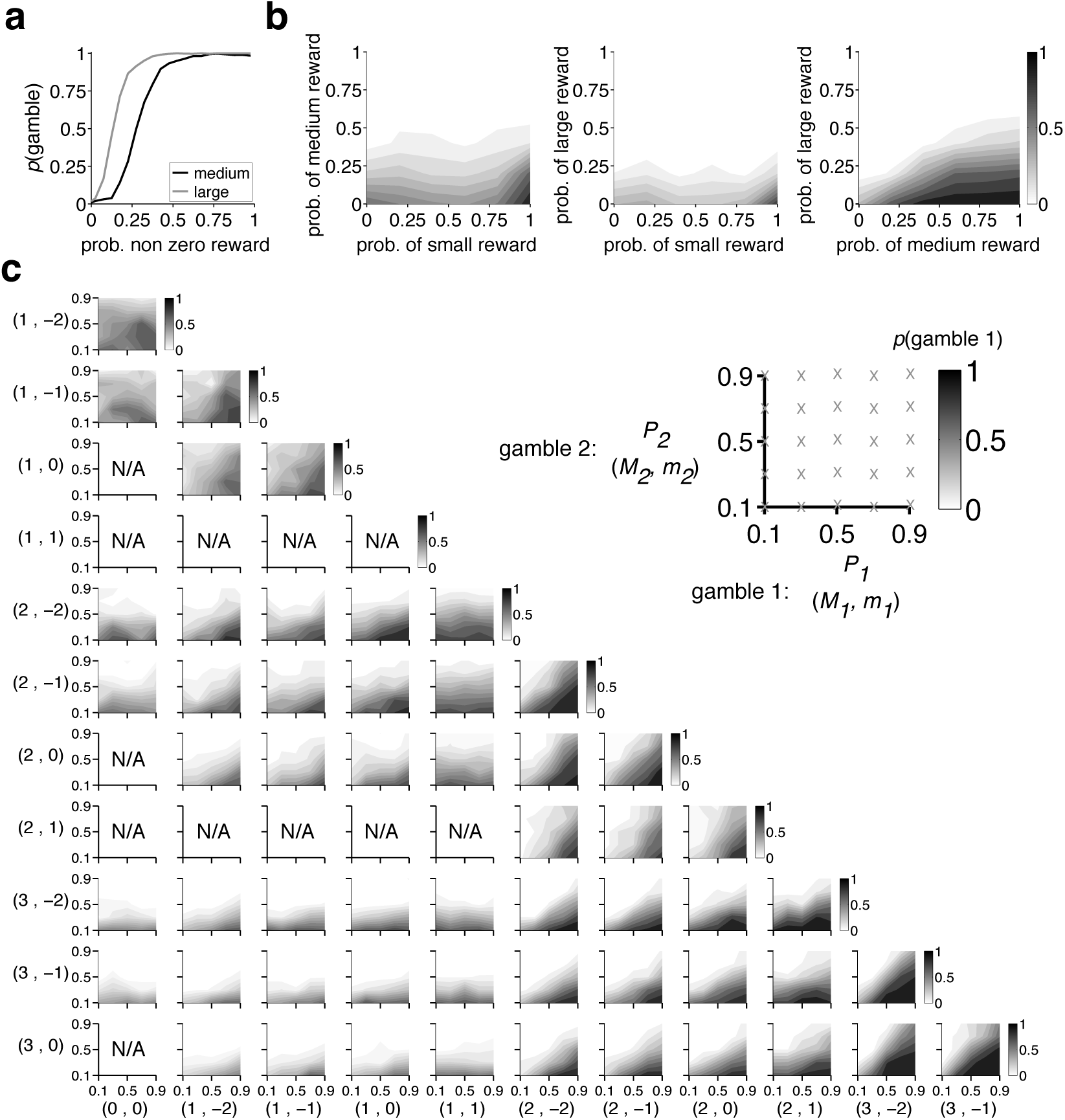
Monkeys’ choices were sensitive to reward probability during both tasks. (**a**) Plotted is the probability of choosing a gamble vs. sure option as a function of the probability of the non-zero reward outcome (medium or large) of the gamble during the juice-gambling task. Selection of the gamble increased monotonically as the probability of non-zero reward outcome increased. (**b**) Each panel plots the probability of choosing a gamble with the non-zero reward outcome indicated on the x-axis as a function of the reward probability of the non-zero outcome of that gamble and the competing gamble (indicated on the y-axis) during the juice-gambling task. (**c**) The same as in (b) for the token-gambling task. Each panel plots the probability of choosing gamble 1 as a function of the reward probability of the larger magnitude outcome of gamble 1 (x-axis) and gamble 2 (y-axis), for a set of gambles with a specific pairs of reward magnitudes. As indicated in the inset, the outcome reward magnitudes of gamble 1 and gamble 2 are shown next to the x- and y-axis, respectively (gamble 1: (*M_1_,P_1_; m_1_,1-P_1_*), gamble 2: (*M_2_,P_2_; m_2_,1-P_2_*)). Overall, the probability of choosing a gamble increased as the probability of reward for its larger magnitude outcome increased, indicating the sensitivity of monkeys to reward probabilities provided by the length of different portions of each bar.

Finally, we examined whether monkeys understood the structure of the token-gambling task and were sensitive to the information about collected tokens that was presented at the bottom of the screen. We reasoned that if monkeys understood this information, then they would necessarily show systematic changes in behavior as a function of token number; for example, they would exhibit more motivation to perform the task as the number of tokens grows and the probability of winning a jackpot reward immediately increases. To test this, we calculated psychometric functions separately for different numbers of collected tokens at the beginning of each trial (**Figure 5**). The psychometric function measures the preference between each pair of gambles as a function of the difference in expected values of gambles, and thus reflects the sensitivity of the animal to the presented information (Eq. 12). We found that both monkeys became less stochastic (smaller *σ* corresponding to a steeper psychometric function) in their decisions, or equivalently more sensitive to the presented information, as they gathered more tokens (*p* = 0.04 for Monkey B and *p* = 0.0003 for Monkey J; two-sided t-test). This result reflects higher level of motivation in performing the task and supports the premise that monkeys can use the token information as a symbolic scoreboard of future rewards.

**Figure 5.**
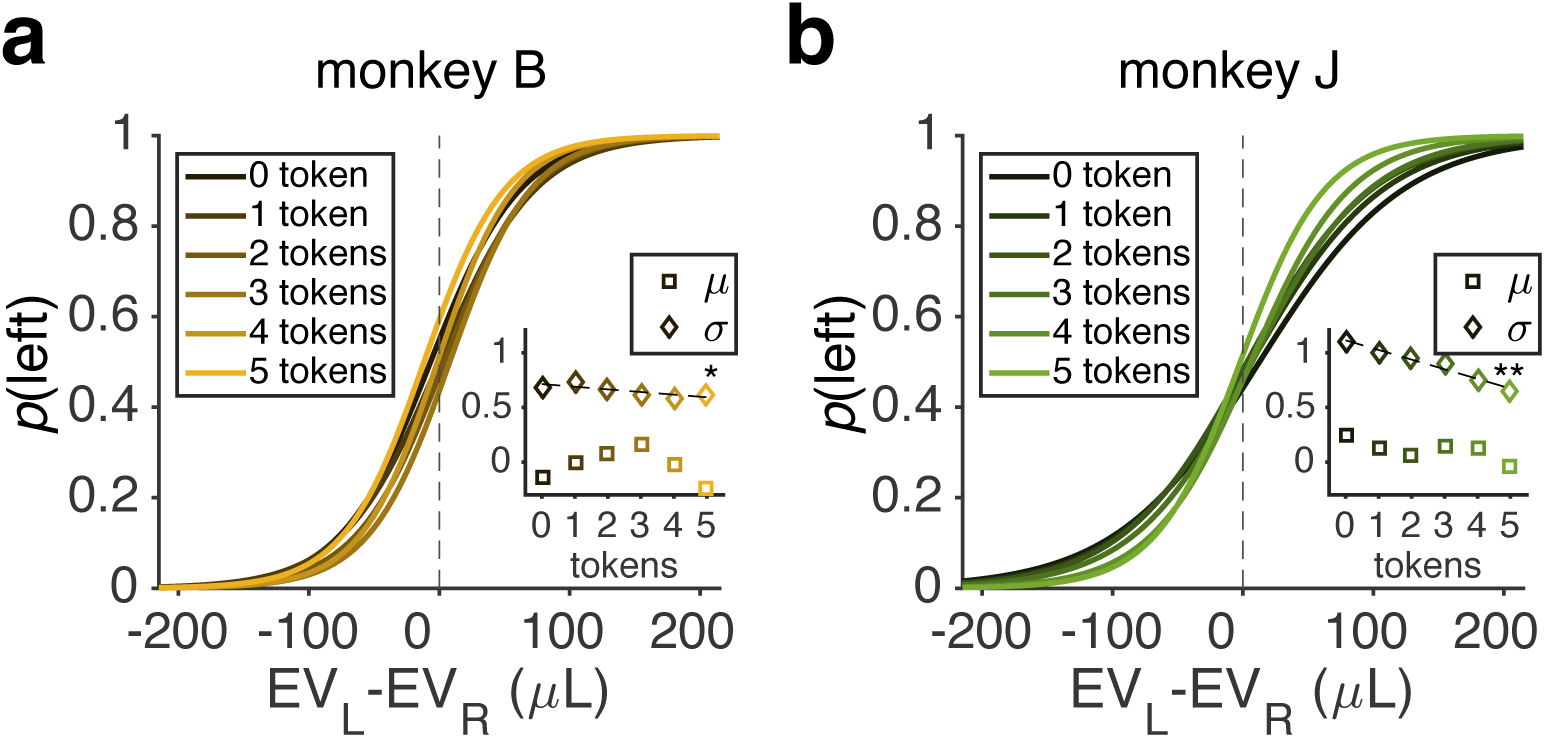
Monkeys’ sensitivity to reward information increased with more collected tokens at the beginning of each trial. **(a)** The probability of choosing the left gamble is plotted as a function of the difference in expected values of two gambles in a given trial, separately for different numbers of collected tokens at the beginning of each trial (shown with different colors). The inset plots the estimated indifference point (*μ*) and stochasticity in choice (*σ*) as a function of different numbers of collected tokens (Eq. 12). One and two stars indicate that the slope of regression line is significantly different from zero at *p* < .05 and *p* < .01, respectively (two-sided t-test). (**b**) The same as (a) but in monkey J. The stochasticity in choice decreased with more collected tokens in both monkeys.

### Monkeys exhibit convex utility curves for both gains and losses and a task-dependent S-shaped probability weighting

Our subjects’ overall risk-seeking and loss-seeking behavior suggests utility and probability weighting functions different from those predicted by prospect theory. More specifically, there are three main characteristics that describe the core risk attitudes of humans in prospect theory (Kahneman and Tversky, 1979). First, the utility curve is concave for gains but convex for losses, indicating risk-aversion and risk-seeking behavior for gains and losses, respectively (**Figure 3a**, blue curve). The opposing risk attitude for gains and losses is known as the reflection effect. Second, the slope of the utility curve for losses is steeper than the one for gains. This pattern produces loss-aversion, the tendency for losses to have a more negative impact on subjective value than equivalent gains. Third, the probability weighting function has an inverse-S-shape, resulting in overweighting of the value of options with small reward probability and underweighting of options with large reward probability (**Figure 3b**, blue curve). To directly assess risk attitudes in monkeys based on prospect theory, we first used four base models to fit choice behavior and estimated utility and probability weighting functions in each of the two experiments (see Materials and Methods). The behavior we observe in our subjects better fits the red curves in **Figures 3a** and **3b**: where a convex curve for gains as well as losses explains risk-seeking behavior in both domains and the probability weighting function (where significant) is S-shaped; suggesting that subjects underweight options with a low probability and overweight options with a high probability. We examine these behavioral patterns in detail below.

Fitting choices based on cross-validation showed that the SU model (the model with nonlinear utility and probability weighting functions; EV: expected value; UF: utility function; PWF: probability weighting function) provided the best fit in the juice-gambling task (**Figure 6a**). Session-by-session estimates of the utility functions based on this model revealed a convex utility function (**Figure 6b**; **Table 1**). This convexity was reflected in the median of the exponent of the utility curve (*ρ_G_*; see Eq. 4 in Materials and Methods) being larger than 1 (median±IQR = ±0.94, two-sided sign-test; *p* < 10^−5^). Monkeys also exhibited a prominent S-shaped probability weighting function (**Figure 6c**) reflected in the distortion parameter (*γ*; see Eq. 5 in Materials and Methods) being larger than 1 (median±IQR = 1.57±0.76, two-sided sign-test; *p* < 10^−5^). Importantly, these results were not model-specific since fitting based on the models with either a nonlinear utility function (EV+UF) or the probability weighting function (EV+PW) also produced convex utility curves (*ρ_G_* median±IQR = 2.57±0.57, two-sided sign-test; *p* < 10^−5^; **Figure 6d**) and a prominent S-shaped probability weighting (*γ* median±IQR = 2.50 ±1.68, two-sided sign-test; *p* < 10^−5^; **Figure 6-1a**).

**Figure 6.**
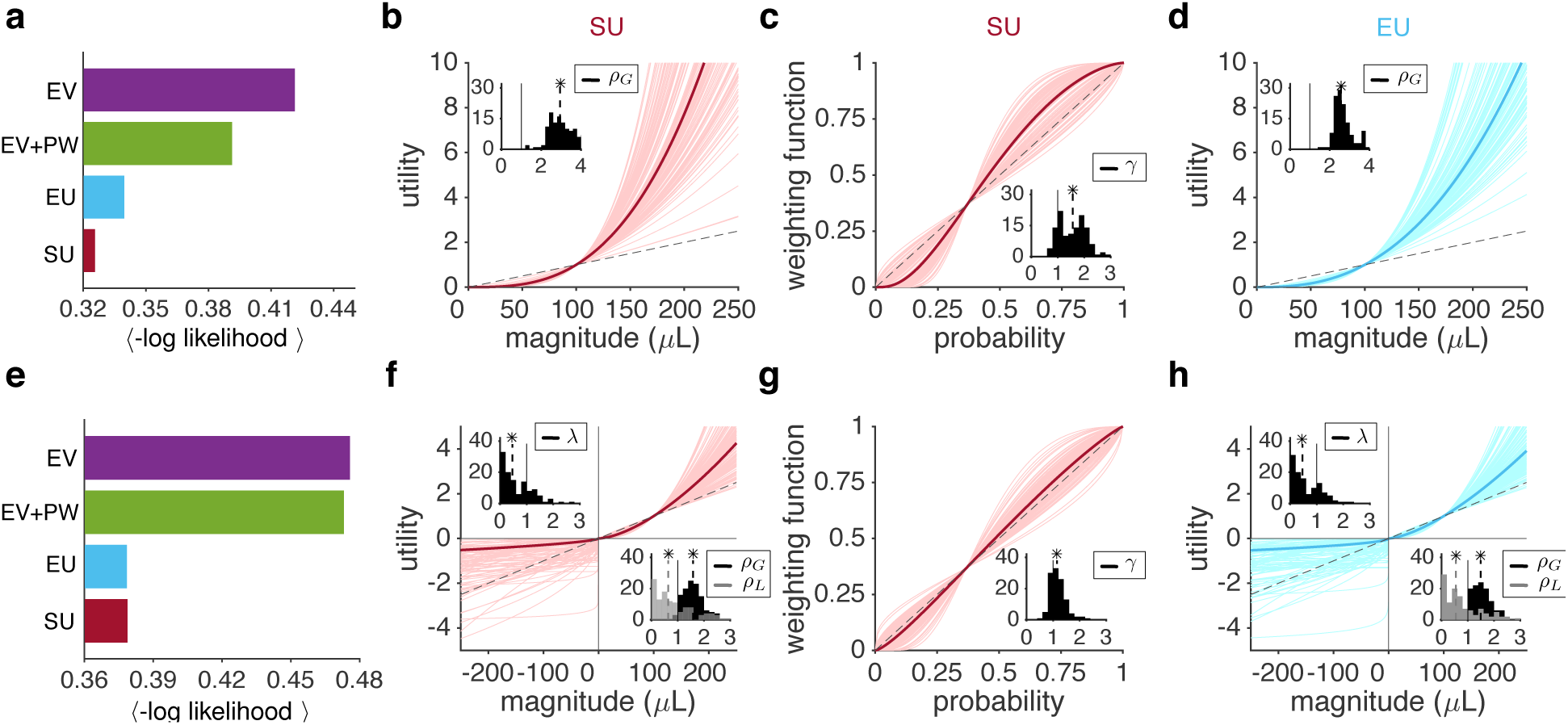
Fitting of choice behavior reveals a task-dependent pattern of risk attitude different than what is predicted by prospect theory. (**a**) Comparison of the goodness-of-fit for choice behavior during the juice-gambling task using four different models of subjective value (EV: expected value; EV+PW: expected value with probability weighting function; EU: expected utility; SU: subjective utility with nonlinear utility and probability weighting functions). Plotted is the average negative log likelihood (−*LL*) per trial over all cross-validation instances (a smaller value corresponds to a better fit). The SU model provided the best fit. (**b**) Estimated utility function based on the SU model as a function of the reward juice. Each curve shows the result of the fit for one session of the experiment; the thick magenta curve is based on the median of the fitting parameter. The dashed line is the unity line normalized by 100*μ*L. The insets show the distribution of estimated parameters for the utility curve for gains (*ρ_G_*). The dashed lines show the medians and a star indicates that the median of the distribution is significantly different from 1 (two-sided sign-test; *p <* 0.05). (**c**) Estimated probability weighting function based on the SU model. The inset shows the distribution of estimated parameters for the probability weighting function (*γ*). The dashed line is the unity line. Subjects exhibited a pronounced S-shaped probability weighting function. **Figure 6-1** shows consistent results for fitting based on the EV+PW model. (**d**) Estimated utility function based on the EU model. Conventions are the same as in (b). (**e-h**) The same as in (a-d), but for the token-gambling task. The EU and SU models provided the best fits and subjects exhibited a slightly S-shaped probability weighting function based on the SU model. The reward magnitude in this task corresponds to the juice equivalent of a given number of tokens. The insets in (f) show the distribution of estimated parameters for the utility curves in the gain (*ρ_G_*) and loss domains (*ρ_L_*), as well as the loss aversion factor (*λ*). On average, subjects exhibited a convex utility function for both gains and losses; they were loss-seeking and violated the reflection effect.

**Table 1.** Summary of the estimated risk preference parameters during the juice-gambling task. Reported are medians (±IQR) of the distribution of estimated parameters in a given model.

|  | SU with cross-opt. DW<br>by mag. | SU | EU | EV+PW |
| --- | --- | --- | --- | --- |
| $\rho_G$ | 1.74±1.00 | 2.95±0.94 | 2.57±0.57 | N/A |
| $\gamma$ | 1.55±0.78 | 1.57±0.76 | N/A | 2.50 ±1.68 |

We next examined choice behavior during the token-gambling task. Fitting choice based on cross-validation showed that the EU and SU models provided the best fit for choice during this task (**Figure 6e**). Fits for the two models were nearly equal suggesting that inclusion of the probability weighting function did not improve the fit and thus the absence of any probability distortion. Session-by-session estimates of the utility functions based on the SU model revealed that monkeys adopted a convex utility function for both gains and losses (**Figure 6f**; **Table 2**). This convexity was reflected in the median of the exponent of the gain utility curve being larger than 1 (median±IQR = 1.58±0.51, two-sided sign-test; *p* < 10^−5^; **Figure 6f** lower inset), and the median of the exponent of the loss utility curve (*ρ_L_*; see Eq. 4 in Materials and Methods) being smaller than 1 (median±IQR = 0.64±1.02, two-sided sign-test; *p* = 10^−4^). In addition, monkeys were loss-seeking: the loss-aversion factor (*λ*; see Eq. 4 in Materials and Methods) was significantly smaller than 1 (median±IQR = 0.46±0.84, two-sided sign-test; *p* < 10^−5^; **Figure 6f** upper inset). Finally, monkeys exhibited a slightly S-shaped probability weighting function (*γ* median±IQR = 1.14±0.37, two-sided sign-test; *p* = 10^−4^; **Figure 6g**). This result is consistent with the finding that the probability weighting function did not improve the fit in the token-gambling task (**Figure 6e**).

**Table 2.** Summary of the estimated risk preference parameters during the token-gambling task. Reported are medians (±IQR) of the distribution of estimated parameters in a given model.

|  | SU with within-opt.<br>DW by mag. | SU | EU | EV+PW | EV |
| --- | --- | --- | --- | --- | --- |
| $\rho_G$ | 1.43±0.46 | 1.58±0.51 | 1.49±0.49 | N/A | N/A |
| $\rho_L$ | 0.48±0.73 | 0.64±1.02 | 0.55±1.05 | N/A | N/A |
| $\lambda$ | 0.58±1.51 | 0.46±0.84 | 0.46±0.84 | N/A | $(\beta_L/\beta_G)$<br>0.13±0.17 |
| $\gamma$ | 1.11±0.4 | 1.14±0.37 | N/A | 1.48±0.58 | N/A |

As with the juice-gambling task, these results were not model-specific: fitting based on the models with either a nonlinear utility function (EU) or the probability weighting function (EV+PW) produced a qualitatively similar pattern of risk preference for the token-gambling task. More specifically, parameter estimates of the utility function based on the EU model showed convex utility curves for both gains and losses that were steeper for the gain than the loss domain (**Figure 6h**; **Table 2**). This was reflected in: the median of *ρ_G_* being larger than 1 (median±IQR = 1.49±0.49, two-sided sign-test; *p* < 10^−5^; **Figure 6h** lower inset); the median of *ρ_L_* being smaller than 1 (median±IQR = 0.55±1.05, two-sided sign-test; *p* = 10^−4^); and the median of *λ* being significantly smaller than 1 (median±QR = 0.46±0.84, two-sided sign-test; *p* < 10^−5^; **Figure 6h** upper inset). Finally, parameter estimates of the probability weighting function based on the EV+PW model revealed an S-shaped weighting function (*γ* median±IQR = 1.48±0.58, two-sided sign-test; *p* < 10^−5^; **Figure 6-1b**). Together, these results show that the observed shape of the estimated utility and probability weighting functions were general and not model-specific.

We also considered the possibility that the observed loss-seeking behavior was caused by monkeys not considering losing a token as a real loss since they were provided with a small ‘motivation’ reward on each trial, regardless of the outcome of the gamble (see Materials and Methods). To test for this possibility, we fit choice behavior with four base models similar to what used above with the difference that a loss of two and one tokens were considered as zero loss or a gain of one token, respectively. The goodness-of-fit based on these models did not reach to those of the models in which losing any token was considered as loss (**Figure 7**). This result strongly suggests that monkeys treated losing tokens as a genuine loss and thus the observed loss-seeking was not due to a shift in the reference point.

**Figure 7.**
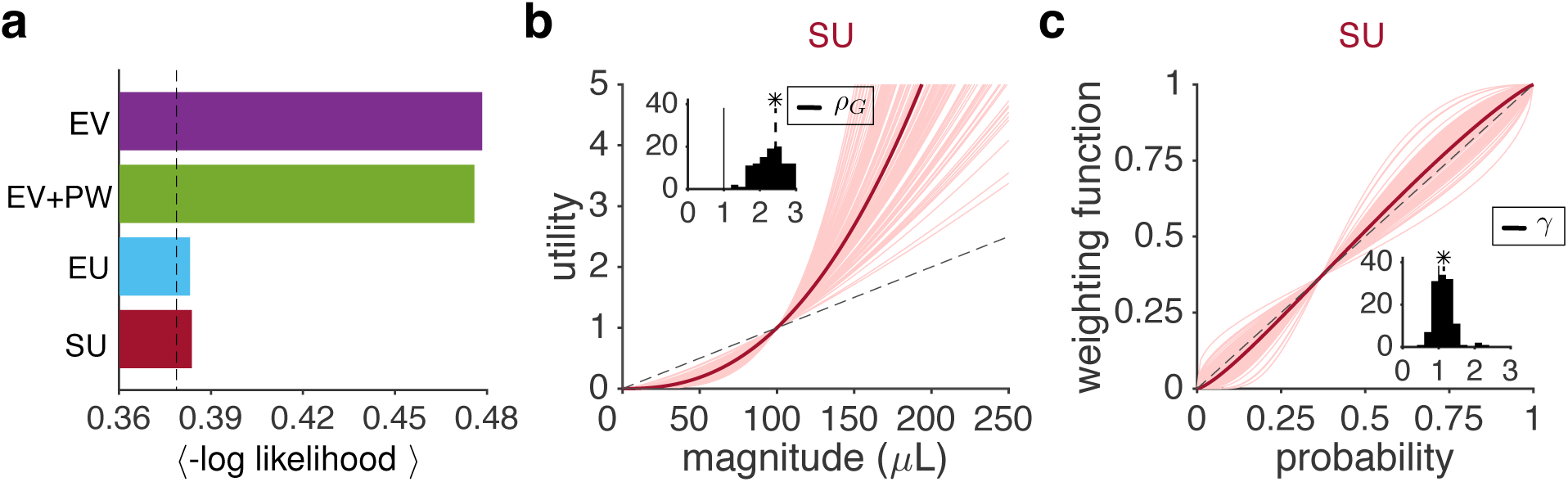
Observed loss-seeking behavior was not due to a shift in the reference point. (**a**) Comparison of the goodness-of-fit for choice behavior during the token-gambling task using four different models of subjective value in which a loss of two and one tokens were considered as zero loss or gain of one, respectively, due to the small ‘motivation’ reward (equivalent to two tokens) provided in each trial. Conventions are the same as in Figure 6. Dashed line indicates the average -*LL* for the best model that considers losing any token as a loss (correspond to the cyan and magenta bars in Figure 6e). The EU model provided the best fit but its goodness-of-fit was worse than the best model that considered losing any token as a loss. (**b**) Estimated utility function based on the SU model as a function of the reward juice. Each curve shows the result of the fit for one session of the experiment; the thick magenta curve is based on the median of the fitting parameter. The dashed line is the unity line normalized by 100*μ*L. The insets show the distribution of estimated parameters for the utility curve for gains (*ρ_G_*). The dashed lines show the medians and a star indicates that the median of the distribution is significantly different from 1 (two-sided sign-test; *p <* 0.05). (**c**) Estimated probability weighting function based on the SU model. The inset shows the distribution of estimated parameters for the probability weighting function (*γ*). The dashed line is the unity line.

Finally, we compared the estimated utility and probability functions in the two experiments. The utility function for gains was significantly more convex in the juice-than token-gambling task (comparison of *ρ_G_* values, two-sided Wilcoxon rank-sum; *p* < 10^−5^). Crucially, this difference was significant even for the two monkeys who performed both experiments (two-sided Wilcoxon rank-sum; *p* < 10^−5^; **Figure 8a, c**). The probability weighting function was more distorted in the juice than token-gambling task (comparison of *γ* values, two-sided Wilcoxon rank-sum; *p* <10^−5^). This pattern held true for both subjects who performed both experiments as well (two-sided Wilcoxon rank-sum; *p* < 10^−5^; **Figure 8b, d**).

**Figure 8.**
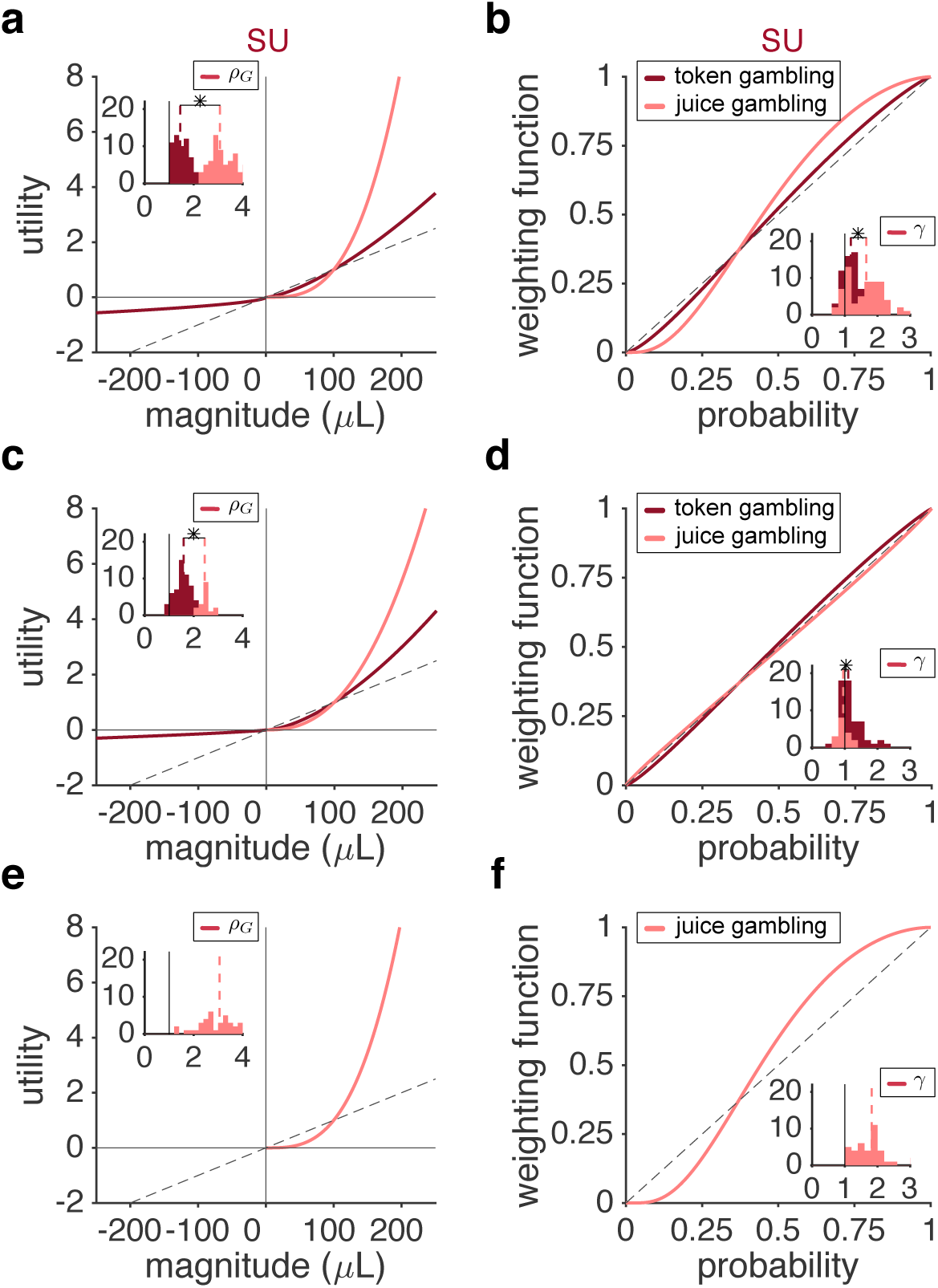
Different utility and probability weighting functions in the two tasks in individual monkeys. (**a**) Estimated utility functions based on the SU model in the juice- and token-gambling tasks. Plot shows the utility curves based on the median of the fitting parameter in the juice (pink) and token (magenta) gambling tasks for monkey B. The dashed line is the unity line normalized by 100*μ*L. The insets show the distributions of estimated parameters for the utility curve for gains (*ρ_G_*) in the two tasks. The dashed lines show the medians, and a star indicates that the medians of the two distributions are significantly different (two-sided Wilcoxon rank-sum; *p <* 0.05). (**b**) Estimated probability weighting function based on the SU model in the two tasks. Plot shows the probability weighting functions based on the median of the fitting parameter in the two tasks for monkey B. The dashed line is the unity line. The insets show the distribution of *γ* values in the two tasks. (**c**-**d**) The same as in (a-b) but for monkey J. Both monkey B and J exhibited a much steeper utility curve in the juice-gambling task. Although both monkeys exhibited different probability distortion in the two tasks this effect was stronger in monkey B. **(e-f**) The same as in (a-b) but for Monkey C that only performed the juice-gambling. The overall behavior of Monkey C was similar to that of Monkey B in the juice-gambling task.

These findings show that risk attitudes, especially in terms of the curvature of the utility function, are flexible and task dependent. To further explore potential mechanisms underlying this flexibility, we next examined additional components involved in the construction of subjective value that could account for some of the observed differences in risk attitudes during the two tasks.

### Differential weighting can partially account for the difference in utility functions across experiments

To explore additional factors that could influence the construction of subjective value and choice, we considered two sets of mechanisms for weighting of possible outcomes. First, we hypothesized that the two gamble outcomes could be weighted differently before they are combined to form the overall subjective value. In other words, the two possible outcomes of a given gamble compete to influence the overall gamble value. To test this hypothesis, we considered three possible ‘within-option’ differential weighting mechanisms by which the gamble outcome with a larger reward magnitude, reward probability, or expected value could influence the overall value more so than the alternative outcome (see Materials and Methods and **Figure 2a-c** for more details). Second, we hypothesized that when comparing two gambles, the value of the better outcome of each gamble could influence its overall value relative to the other gamble. To test this hypothesis, we considered alternative ‘cross-option’ differential weighting mechanisms based on the magnitude, probability, or expected value of the better outcome in each gamble (see Materials and Methods and **Figure 2d-i** for more details). We used all these models to fit choice behavior in the two experiments.

We found that the SU model with cross-option differential weighting (DW) based on reward magnitude provided the best fit in the juice-gambling task (**Figure 9a**). In order to study the contribution of DW to flexible risk attitudes, we next compared the session-by-session estimates of the ‘differential-weighting factor’ (see Materials and Methods) and risk preference parameters based on the SU model with and without DW. This analysis revealed a strong differential weighting of the two gambles based on reward magnitude of the better outcome (DW factor median±IQR = 0.63±0.18, two-sided sign-test; *p* < 10^−5^; **Figure 9d**) corresponding to ∼102% larger weight for the value of the gamble with the larger magnitude relative to the other gamble. More importantly, the estimated utility function was less steep in the SU model with differential weighting than in the SU model without differential weighting (*ρ_G_* median±IQR = 1.74±1.00 and 2.95±0.94 for the model with and without DW, respectively; two-sided sign-test, *p* < 10^−5^; **Figure 9b; Figure 10a**). This finding suggests that differential weighting accounts for some portion of behavior that, unless modeled explicitly, influences estimates of the subjective utility function, suggesting stronger value distortions. However, there was no difference between probability distortion estimates based on the model with and without DW (*γ* median±IQR = 1.55±0.78 and 1.57±0.76 for the model with and without DW, respectively; two-sided sign-test, *p* = 0.12; **Figure 9c; Figure 10b**), suggesting that differential weighting may not influence estimates of this function, at least not in this task.

**Figure 9.**
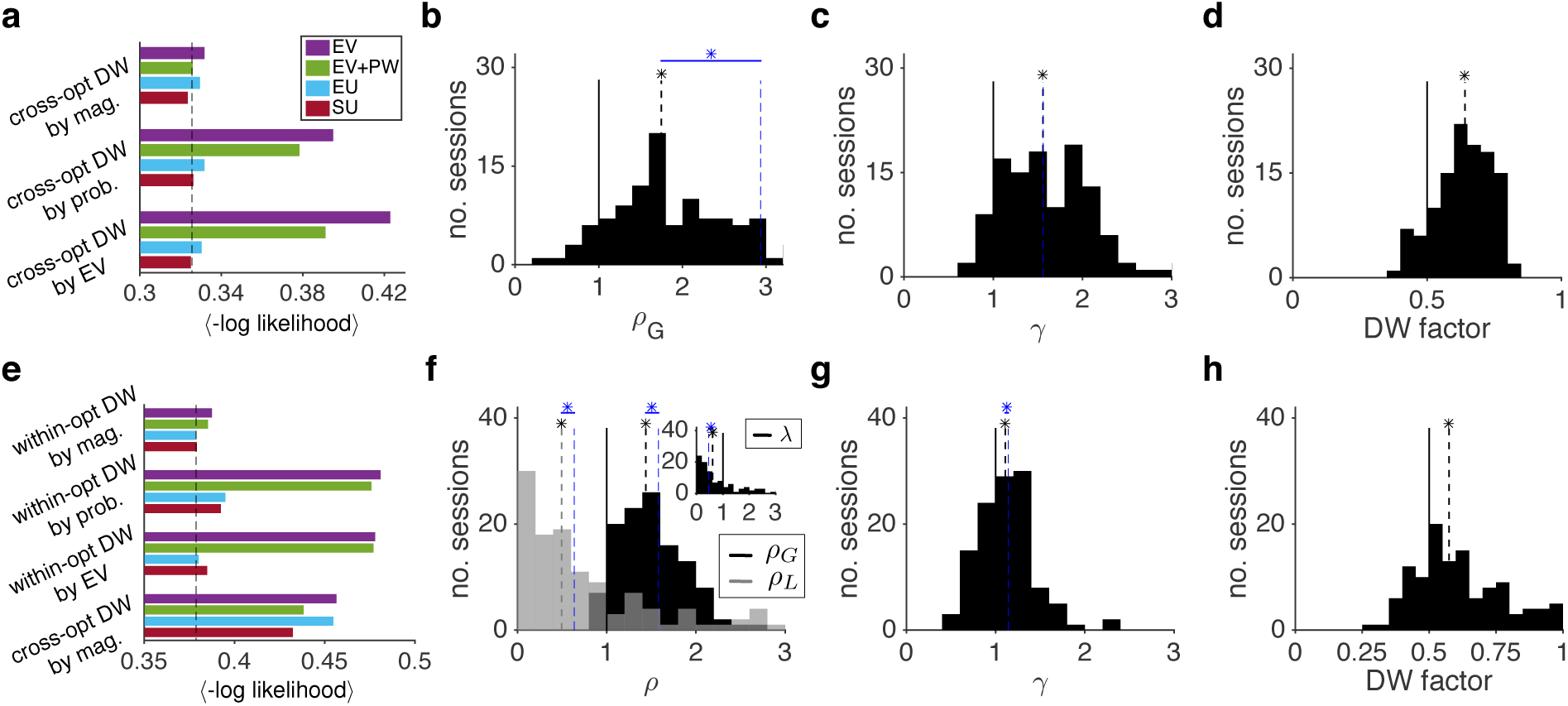
Differential weighting of gamble outcomes can account for part of the convexity of the utility functions. (**a**) Comparison of the goodness-of-fit for choice behavior during the juice-gambling task using four different models for the construction of reward value for each possible outcome (convention the same as in Figure 6) and three different types of differential weighting (DW) mechanisms (cross-opt: cross-option). Plotted is the average negative log likelihood (−*LL*) per trial over all cross-validation instances (a smaller value corresponds to a better fit). Overall, the SU model with cross-option DW by magnitude provided the best fit in the juice-gambling task. Dashed line indicates the average -*LL* for the best model without DW. (**b-c**) Distributions of the estimated parameters for the utility (b) and probability weighting function (c) using the SU model with DW. The black dashed lines show the medians, and a black star indicates that the median of a given distribution is significantly different than 1.0 in (b-c) and 0.5 in (d) (two-sided sign-test; *p <* 0.05). The blue dashed line shows the median of the best model without DW (the same medians as in **Figure 6b-c**). A blue star indicates that the estimated parameter was significantly different between the best models with and without DW (two-sided sign-rank test; *p <* 0.05). Differential weighting can account for part of the convexity of the utility function since the model with this mechanism is less convex. (**d**) Distribution of estimated DW factors using the SU model with DW. There was a significant DW effect in favor of the gamble with the larger reward magnitude. (**e-h**) The same as in (a-d), but for the token-gambling task. Overall, the EU and SU with within-option DW by magnitude models provided the best fit. Moreover, all models with cross-option DW by magnitude were provided worse fits than corresponding models with within-option DW.

**Figure 10.**
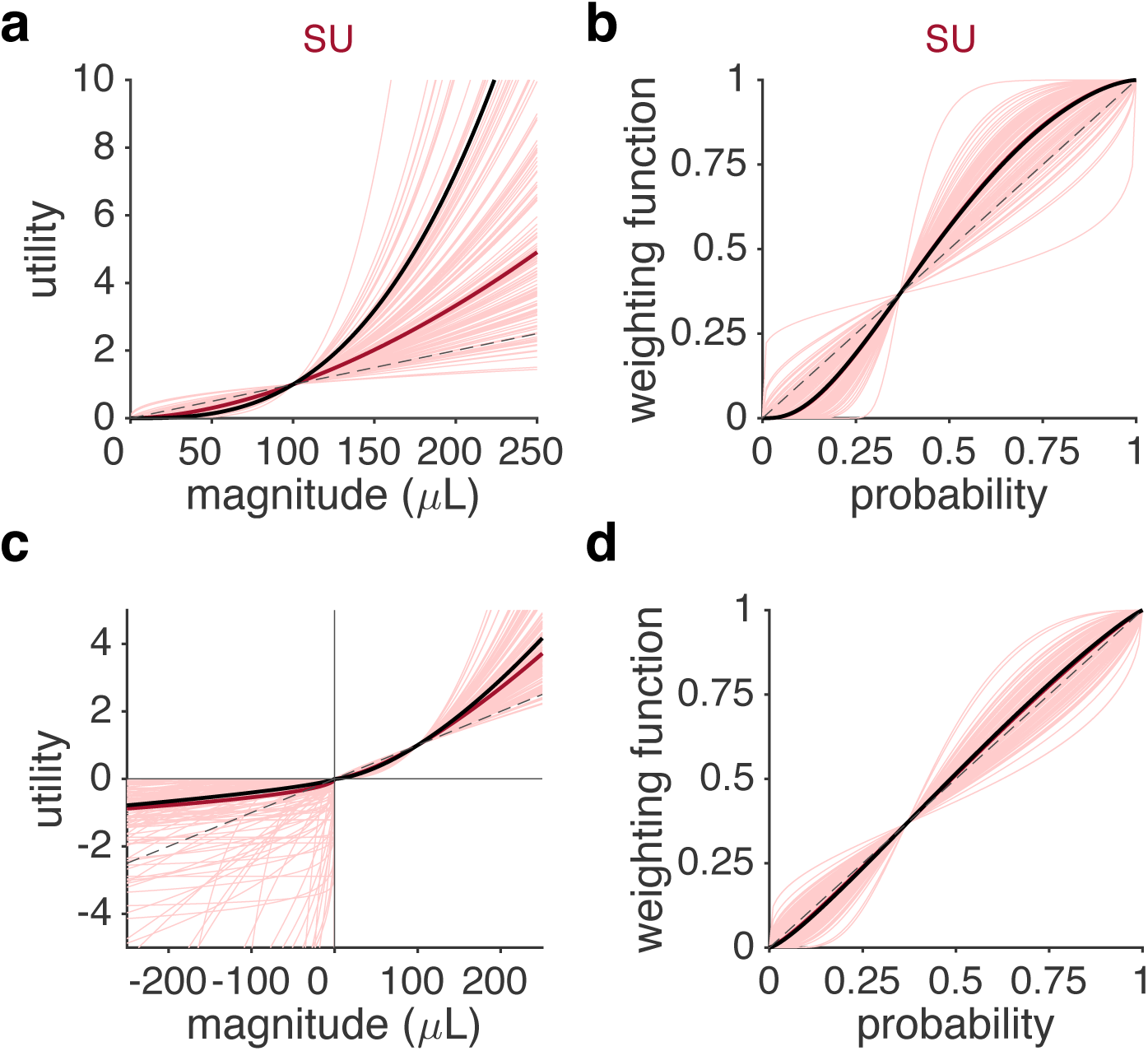
Estimated utility and probability weighting functions for the SU model with differential weighting by reward magnitude. (**a-b**) Estimated (a) utility and (b) probability weighting functions based on the SU model with within-option differential weighting by magnitude for the juice-gambling task. Each curve shows the result of the fit for one session of the experiment, and the thick red curve is based on the median of the fitting parameter. For comparison, the black curves show the utility (a) and probability weighting functions based on the SU model without DW. The probability weighting curves are indistinguishable in the two models. The dashed line is the unity line normalized by 100 *μ*L in (a), and the unity line in (b). (**c-d**) The same as in (a-b) but for the token-gambling task.

In contrast to the juice-gambling task, models with within-option DW provided better fit compared to models with cross-option DW in the token-gambling task (compare bottom and top four bars in **Figure 9e**). Overall, the EU and SU models with within-option DW based on reward magnitude provided the best fit among all models with DW. The improvement of fit based on theses models relative to the models without DW was minimal (base EU and base SU models, **Figure 6e**). These results indicate that differential weighting did not strongly influence choice behavior in the token-gambling task. Nevertheless, the session-by-session estimate of the DW factor in the SU with DW model revealed a significant effect of DW on valuation; DW factors were significantly larger than 0.5 (median±IQR = 0.57±0.22; two-sided sign-test; *p* < 10^−5^; **Figure 9h**) corresponding to ∼33% larger weight for the value of the outcome with the larger magnitude relative to the outcome with the smaller reward magnitude.

Moreover, the utility functions for both gains and losses were less steep in the SU model with DW than in the SU model without DW (**Figure 9f; Figure 10c**; **Table 2**). The estimated exponents of the utility function for gains (*ρ_G_*) were significantly smaller after considering differential weighting (median±IQR = 1.43±0.46 and 1.58±0.51 for the model with and without DW, respectively; two-sided sign-test, *p* = 1.3*10^−4^). Similarly, the estimated exponents of the utility function for losses (*ρ_L_*) were significantly smaller after considering differential weighting (median±IQR = 0.48±0.73 and 0.64±1.02 for the model with and without DW, respectively; two-sided sign-test, *p* = 0.048). The estimated loss aversion factors, however, were larger in the model with DW (*λ* median±IQR = 0.58±1.51 and 0.46±0.84 for the model with and without DW, respectively; two-sided sign-test; *p* < 10^−5^) corresponding to more loss-seeking in this model. Finally, the probability weighting function was slightly less distorted in the model with differential weighting (*γ* median±IQR = 1.11±0.40 and 1.14±0.37 for the model with and without DW, respectively; two-sided sign-test; *p* < 10^−5^; **Figure 9g; Figure 10d**). These results demonstrate that within-option differential weighting can account for some of the observed convexity of the utility functions in the token-gambling task.

To demonstrate that our fitting procedure can actually distinguish between alternative models and identify the correct model and to accurately estimate model parameters, we generated choice data using all the presented models and over a wide range of model parameters, and subsequently fit the simulated data with all the models (see Materials and Methods). We found that data generated with certain models were easier to fit than other models. For example, models without DW were in general easier to fit, and within a given family of models, data generated with models with non-linear utility functions (EU and SU) were easier to fit (**Figure 2-1a,c**). Nevertheless, the same model used to generate a given set of data provided the best overall fit (**Figure 2-1b,d**). We also computed the relative estimation error (i.e. difference between the estimated and actual parameters after normalizing each estimated parameter by its actual value; see Materials and Methods) and found that fitting based on the model used to generate a given set of data provided an unbiased estimate of model parameters (**Figure 2-2a,c**). Moreover, we found the minimum value of the average absolute estimation error (as a more robust measure of variance in estimation error) for the same model used to generate a given set of data (**Figure 2-2b,d**). Together, these results demonstrate that our fitting method is able to correctly identify the model used to generate a given set of data and thus can distinguish between the alternative models. Moreover, our fitting yields unbiased estimates of model parameters with relatively small error.

Altogether, these results illustrate that differential weighting could account for part of the observed convexity of the utility function in both experiments. Interestingly, the amount of change in the convexity after including DW was larger in the juice-gambling task (Δ*ρ_G_* median±IQR = −1.09±1.52) than in the token-gambling task (Δ*ρ_G_* median±IQR = −0.09±0.20; two-sided Wilcoxon rank-sum test, *p* < 10^−5^), making the utility functions more similar after the inclusion of DW (**Figure 11a**). We did not observe similar changes in the estimates of probability distortion parameters after the inclusion of DW (juice task Δ*γ* median±IQR = 0.02±0.13; token task Δ*γ* median±IQR = −0.03±0.10; two-sided Wilcoxon rank-sum test, *p* = 0.27; **Figure 11b**). These results demonstrate that the differential weighting mechanisms can partially account for the observed difference in utility function across the two tasks. Moreover, they explain how such additional mechanisms enable flexible risk attitude according to the task.

**Figure 11.**
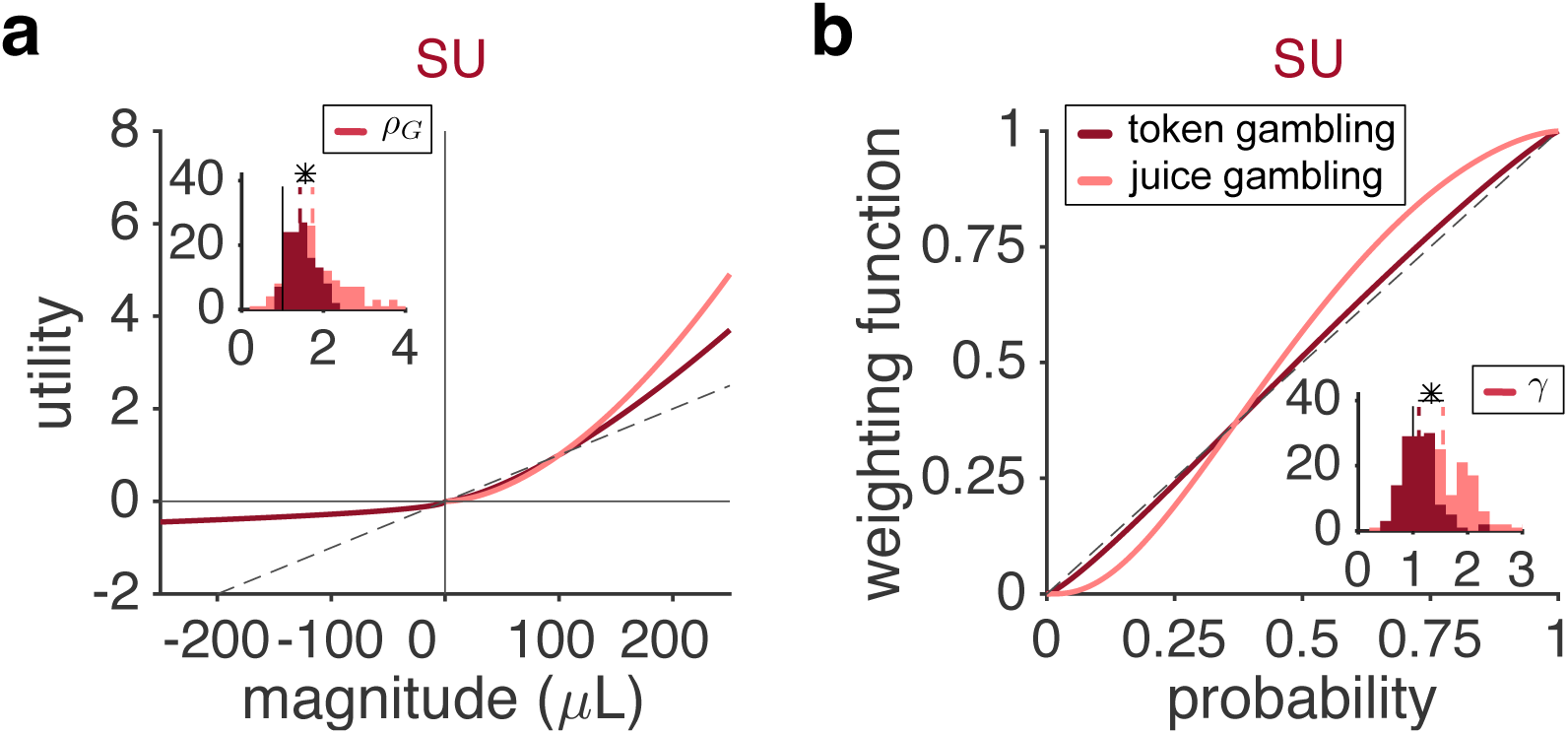
Comparison of the estimated utility and probability weighting functions in the juice- and token-gambling tasks based on the SU model with differential weighting. (**a**) Estimated utility functions based on the SU model with differential weighting in the juice- and token-gambling tasks. Plot shows the utility curves based on the median of the fitting parameter in the juice (pink) and token (magenta) gambling tasks. The dashed line is the unity line normalized by 100*μ*L. The insets show the distributions of estimated parameters for the utility curve for gains (*ρ_G_*) in the two tasks. The dashed lines show the medians. There was a small but significant difference between the two distributions (two-sided Wilcoxon rank-sum; *p <* 0.05). (**b**) Estimated probability weighting function based on the SU model with differential weighting in the juice- and token-gambling tasks. Plot shows the probability weighting functions based on the median of the fitting parameter in the two tasks. The dashed line is the unity line. Other conventions are the same as in panel (a). The star indicates that the medians of the two distributions are significantly different (two-sided Wilcoxon rank-sum; *p <* 0.05). Correlations between estimated model parameters are plotted in **Figures 11-1**, **11-2**, and errors in the estimation of model parameters are shown in **Figure 11-3**.

We also calculated the correlations between the estimated parameters of the best models (the SU models with and without DW) in order to test whether some of the observed effects of differential weighting could be captured by changes in other parameters. We calculated these correlations using two different methods: the Hessian matrix and session-by-session values of fitting parameters (see Materials and Methods). These analyses revealed that in both models the exponent of utility function power law (*ρ_G_*) and the stochasticity in choice (σ) were significantly correlated with each other, indicating that a larger amplification of reward magnitude by the utility function can be offset with a larger value for stochasticity in choice (**Figure 11-1** and **Figure 11-2**). This correlation can be an evidence for normalization in value construction. Additionally, we found that in the SU model with differential weighting, differential-weighting factor (*DW*) is significantly correlated with ρ_G_ and σ. Moreover, in the juice-gambling task, the *DW* parameter was significantly correlated with *ρ_G_* and to a lower extent with *σ*. In the token-gambling task, we also found a correlation between *DW* and *σ*, and between *DW* and loss aversion factor (*λ*). It is worth noting that the observed correlations should not be concerning for the interpretation of best fitting models because we used cross-validation for identifying those models. Cross-validation would reveal if any of the fitting parameters in our best models were redundant.

One possible concern could be that because of the correlation between *ρ* and *σ*, some of the observed change in *ρ* (i.e. the convexity of the utility function) between the two tasks could be caused by changes in *σ* and not differential weighting. To rule out such possibility, we defined a single quantity for measuring the effect of reward magnitude on choice behavior equal to *x*^*ρ*^*_G_*/*σ* (and *λx*^*ρ*^*_L_*/*σ* for losses), where *x* is one of the possible reward magnitudes. The value of *x*^*ρ*^*_G_*/*σ* determines the influence of reward magnitudes on choice in a given model considering the stochasticity in choice in that model. We then computed the distributions of *x*^*ρ*^*_G_*/*σ* for the best models with and without DW and found that *x*^*ρ*^*_G_*/*σ* (and *λx*^*ρ*^*_L_*/*σ*) values were significantly larger in the SU with DW model (two-sided Wilcoxon rank-sum; *p <* 0.05; **Figure 12**). These results show that despite correlations between *DW* and *ρ_G_* and *σ*, differential weighting results in enhanced value of reward magnitude relative to stochasticity in choice. This indicates that differential weighting of possible outcomes based on the magnitude increases the overall effect of magnitude on choice and thus can capture some of risk-seeking behavior that are otherwise attributed to the convexity of the utility function.

**Figure 12.**
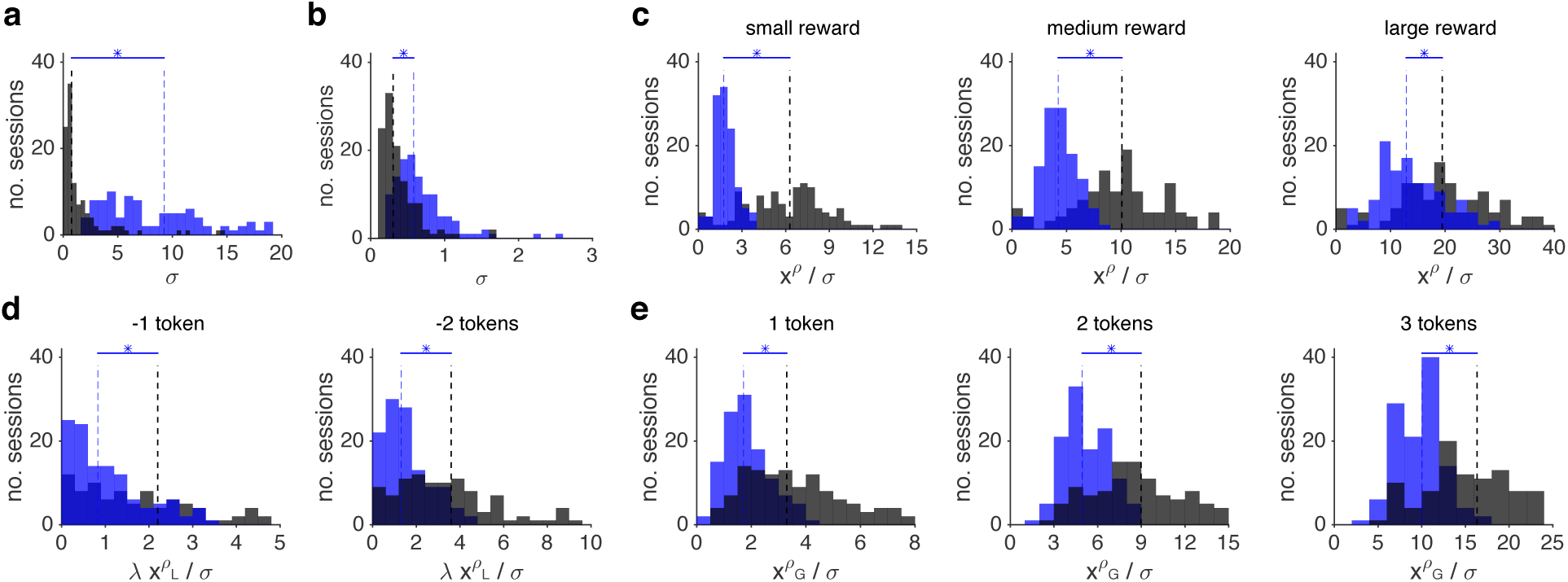
Differential weighting of possible outcomes enhances the overall effect of magnitude on choice. (**a-b**) Distribution of estimated values of stochasticity in choice (*σ*) during the juice-gambling task (a) and the token-gambling task (b). The SU model with and without DW are shown in blue and black, respectively. The dashed lines show medians, and a blue star indicates that the medians of two distributions are significantly different from each other (two-sided Wilcoxon rank-sum; *p <* 0.05). Stochasticity in choice was significantly smaller in the SU with DW model. (**c**) Distributions of *x*^*ρ*^/*σ* values for small, medium, and larger rewards during juice-gambling task. (**d-e**) Distribution of *λx*^*ρ*^*_L_*/*σ* values for loss tokens (d) and *x*^*ρ*^*_G_*/*σ* values for gain tokens (e) during the token-gambling task. In all cases, the *x*^*ρ*^/*σ* values were significantly larger in the SU with DW model indicating that differential weighting of possible outcomes based on the magnitude increases the overall effect of magnitude on choice.

Finally, we also examined the likelihood surface of the best models in order to calculate the error associated with the estimated parameters. Overall, we found small errors in the estimation of model parameters, expect for a few parameters that were correlated with other parameters: *σ* in the SU with DW model in the juice-gambling task, *ρ_L_* in the SU model in the token-gambling task, and *ρ_L_* and *λ* in the SU with DW model in the token-gambling task (**Figure 11-3**).

Importantly, estimation errors in these parameters do not affect our results.

## DISCUSSION

### Flexible risk attitudes in monkeys

We investigated risky choices in monkeys performing two different gambling tasks: a token-gambling task (with both gains and losses) and a juice-gambling task (with gains only). Fitting choice behavior with alternative models revealed convex utility curves for both the gain and loss domains, a pattern that is inconsistent with the reflection effect. Macaques thus deviated from humans and capuchins (Lakshminarayanan et al., 2008). Moreover, our monkeys showed a steeper utility curve for gains than for losses, making them loss-seeking; a deviation from the loss-aversion observed in humans and capuchins (Chen et al., 2006). Finally, monkeys showed a prominent S-shaped probability weighting fucntion in the juice-gambling task and nearly linear (albeit slightly S-shaped) probability weighting in the token-gambling task; these patterns deviate from each other, from previous human studies, and from rhesus macaques in two other studies (Stauffer et al., 2015; Yamada et al., 2013).

Taken together, our results challenge the idea that rhesus monkeys have a fixed and stable set of risk attitudes that are consistent across tasks. This variety in responses to risk challenges the idea that these risk attitudes have not changed since the time of our most recent ancestor. Instead, our results support an alternative view in which natural selection in the primate order has led to robust cognitive flexibility. This flexibility, presumably, would prevent us from having risk attitudes that are so ingrained that we would fail to rapidly adjust our utility curves or probability weighting to changing task conditions. In contrast, the flexibility requires mechanisms that can be adjusted to the task at hand; for example, different utility and probability weighting functions for different tasks.

### Neural mechanisms of flexibility in risk attitudes

A major goal of neuroeconomics has been to understand how our responses to uncertainty are determined by, presumably, specially dedicated neural mechanisms. It is often assumed that risk attitudes are stable and that the goal of neuroeconomics then is to understand how relevant neural operations lead to these preferences. Our work points to a different possibility: if preferences are not stable, then the neural processes that produce them may be similarly flexible. Indeed, our results suggest a somewhat different desideratum: that neuroeconomics ought to focus on how the brain regulates risk attitudes in response to context and adjusts them rapidly and adaptively when demands change. More broadly, and more speculatively, our findings suggest that risk attitudes may be seen as a consequence of general neural mechanisms that support rapid adjustment, presumably in contexts divorced from risk, rather than of a special and dedicated uncertainty module in the brain. Our results point to attentional modulation as a plausible mechanism. All these results are relevant for future studies into neural mechanisms of value computations and how they are adjusted.

Standard approaches to modeling choice, especially in the neural domain, hold that different prospective outcomes of a single offer are weighted equally in evaluation (that is, after all, the normative strategy as well as the simpler one). It is surprising then that our results point to two types of differential weighting based on reward magnitude: a within-option weighting for outcomes within a risky option and an cross-option weighting for the two options. These findings can be explained by the idea that the weighting processes that determine value are biased by the greater attentional weight assigned to some prospects (typically the more salient outcomes, Hayden et al., 2008; Hayden and Platt, 2007; Ludvig et al., 2014; Shimojo et al., 2003; Busemeyer & Townsend, 1993; Roe, Busemeyer, & Townsend, 2001). In the juice-gambling task in which there is only one non-zero outcome per gamble, competitive differential weighting occurs between the two gambles –perhaps via spatial attention. In the token-gambling task in which there could be two non-zero outcomes in each gamble, the competitive differential weighting occurs within a gamble –perhaps via feature-based attention. Even though models with a differential weighting mechanism only minimally improved the quality of fit in this task, the result of the comparison of fitting parameters indicates that this mechanism can account for part of the convexity of the utility function. Our results then illustrate how attentional mechanisms can influence economic decisions and make them more flexible. Empirically, modeling the influence of attention on evaluation is essential because some of the variance attributed to utility curves may actually reflect reweighting instead. Traditional approaches that do not take this possibility into account may over-estimate the curvature of the utility function.

### Stable vs. constructed values and comparison with previous studies

One tradition in behavioral economics holds that preferences are *constructed* at the time of elicitation, and do not *reveal* stable values (Lichtenstein and Slovic, 2006). The set of computations involved in preference construction include not only a value function, but also editing, reference dependence, reweighting, and so on. Our results appear to be consistent with this view. These patterns are not likely to be restricted to the domain of risk; they are also consistent, for example, with an emerging body of work showing that preferences in the time domain are highly dependent on seemingly irrelevant contextual factors (Stephens and Anderson, 2001; Pearson, Hayden, and Platt, 2010; Blanchard and Hayden, 2015; reviewed in Hayden, 2016). Together, these results suggest that both risky and temporal preferences are constructed in animals, and thus extend the concept of preference construction beyond humans.

We have previously argued that, when playing fast repeated gambles for small amounts, monkeys are more likely to focus on the win than on the loss (Hayden and Platt, 2007; Hayden, Heilbronner, Platt, 2009), and that humans may do the same when confronted with those contexts (Hayden and Platt, 2009). Our results here provide three pieces of evidence for this argument. First, a convex utility curve that became steeper from losses to gains showed how larger wins were valued more. Second, we found a strong differential weighting based on reward magnitude across gambles in the juice-gambling task, indicating that, indeed, a larger win strongly influenced the behavior. Third, in the task with both gains and losses (token-gambling task), monkeys again differentially weighted possible outcomes of each gamble based on reward magnitude.

One factor that could explain the shape of the probability weighting function is the difference between description and experience in communicating the properties of the gamble (Hertwig et al., 2004; Hertwig and Erev, 2009; Ludvig and Spetch, 2011). Humans, like our monkeys, exhibit S-shaped curves in experienced gambles. It may be that monkeys in our tasks treated the gambles as more described-like than experienced-like, especially in the juice-gambling task in which we used a much higher resolution for reward probability (0.02 vs. 0.2 in the juice vs. token task, respectively). Monkeys could trust a larger set of reward probabilities less and therefore rely more on experience to evaluate corresponding gambles. Reliance on experience is a useful strategy for tackling reward uncertainty (Farashahi et al., 2017). In a recent study showing an inverse-S-shape for probability distortion in monkeys, only six values of reward probability were used, which could have made the gambles act as more described (Stauffer et al., 2016). Note also that, in that study, only one value for the reward magnitude was used in gambles. This limitation could result in degeneracy in fitting solutions, causing the inverse-S-shaped probability weighting to absorb some the convexity of the utility function.

Ultimately, these results serve as a testament to the cognitive flexibility and adaptiveness of rhesus monkeys, which are among the most successful primate species (Strier, 2016). Indeed, the success of the rhesus macaque is in part attributable to its ability to adjust to changing environments, including an omnivorous diet and a willingness to live in a variety of climates, ranging from warm to cold as well as locations in both city and country. That is, regardless of the dietary richness of the environment in which they evolved, rhesus monkeys have thrived because they can adjust to new environments rapidly. Thus, in our view, it should not be surprising that they do not have a stable set of risk attitudes. It remains an open question how these ideas relate to species with narrower niches.

## Acknowledgments

We would like to thank Rei Akaishi, Ben Eisenreich, Clara Guo, Daeyeol Lee, and Katherine Rowe for helpful comments on the manuscript. This work is supported by a CAREER award from NSF (BCS1253576) and a R01 from NIH (DA038615) to BYH, and by NSF EPSCoR (RII Track-2 FEC) grant to AS. We thank Meghan Castagno, Marc Mancarella and Caleb Strait for assistance with data collection.

---

**Figure 2-1.**
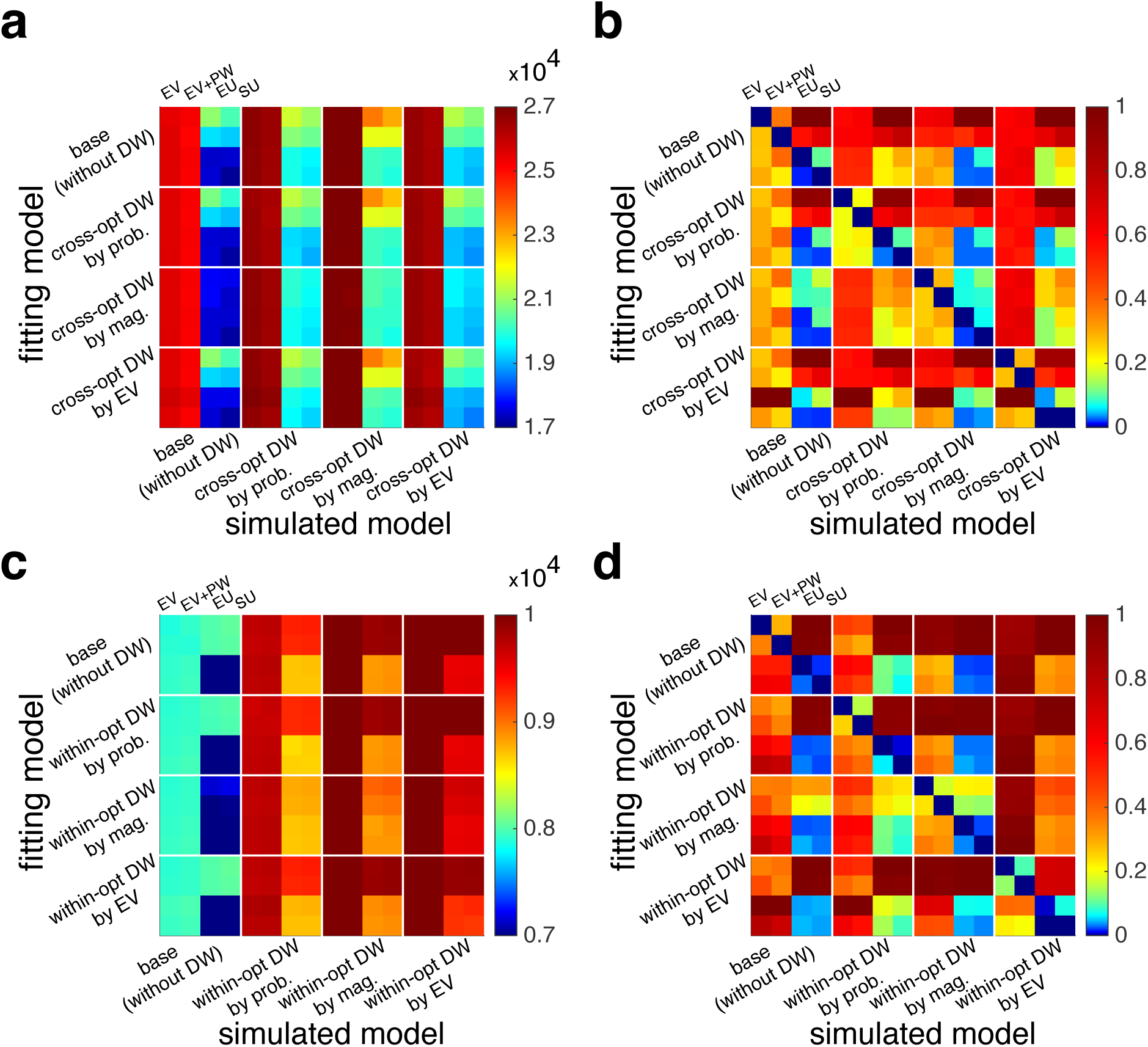
Fitting procedure is able to identify the model used for simulating data correctly. (**a**) Plot shows the goodness-of-fit (in terms of the average AIC over a set parameters) for fitting choice data generated with a given model and fit with the same or different models (total 16 models) for the juice-gambling task. The models used to generate and fit data are indicated on the x- and y-axis, respectively. (**b**) The same as in (a), but the AIC values for data generated with a given model (each column) are rescaled by first subtracting the minimum AIC value obtained by fitting a given set of data and then dividing the outcome by the difference between the maximum and minimum values of AIC for that set of data. As a result, rescaled AIC values for each set of simulated data fall between 0 and 1. (**c-d)** The same as in (a-b) but for the token-gambling task. Overall, the same model used to generate a given set of data provided the best overall fit.

**Figure 2-2.**
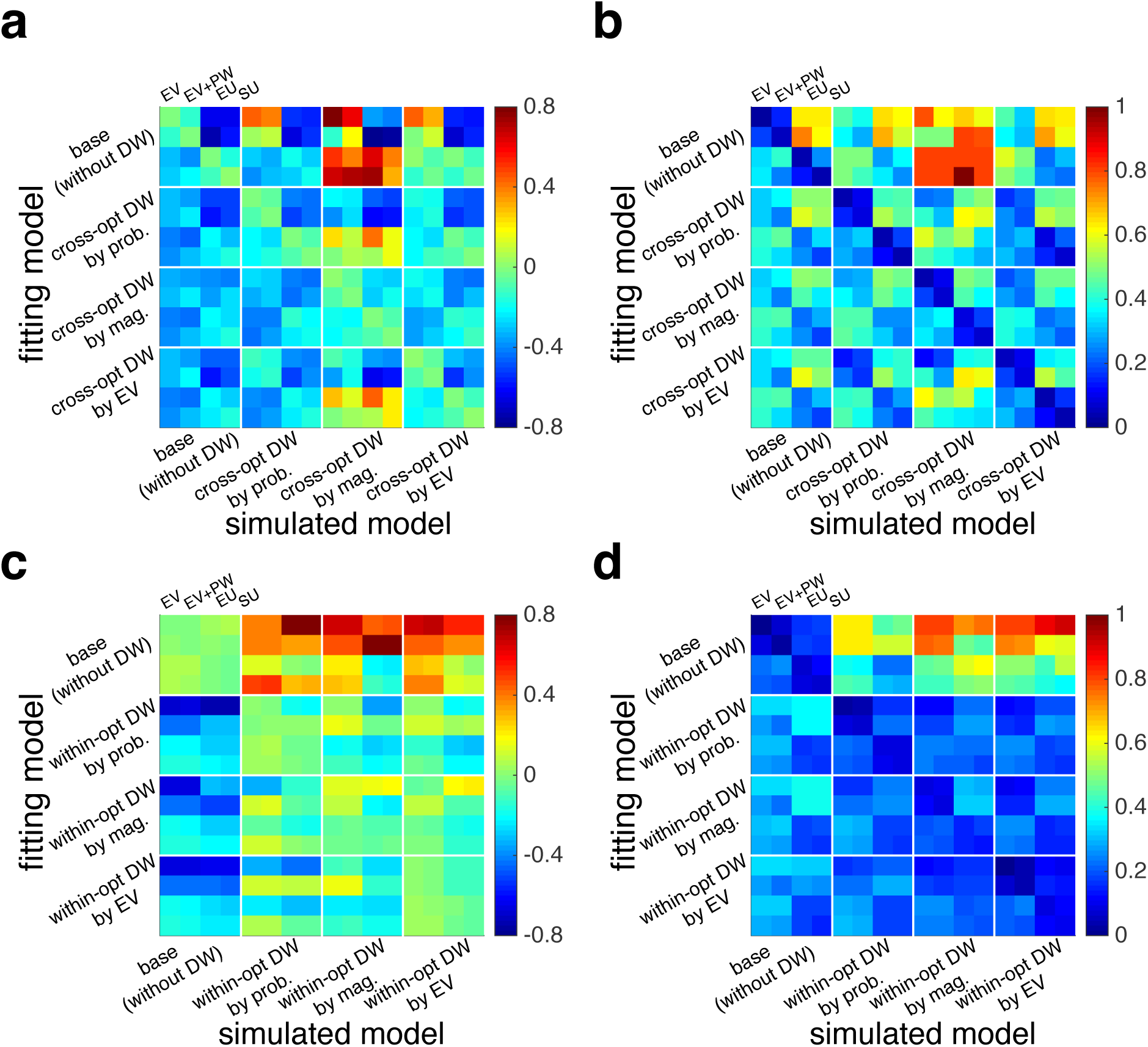
Fitting procedure is able to estimate model parameters accurately and with relatively small error. **(a)** Plotted is the average relative estimation error (i.e. the difference between the estimated and actual parameter values divided by the actual value) in fitting choice data generated with a given model and fit with the same or different models (total 16 models) for the juice-gambling task. The fit using the same model used to generate a given set of data provided unbiased estimates of model parameters. (**b**) Plotted is average of the absolute value of relative estimation error (as a more robust measure of variance in estimation error) in fitting choice data generated with a given model and fit with the same or different models. The variance in estimated parameters was the minimum for fit using the same model used to generate a given set of data. (**c-d**) The same as in (a-b) but for the token-gambling task.

**Figure 6-1.**
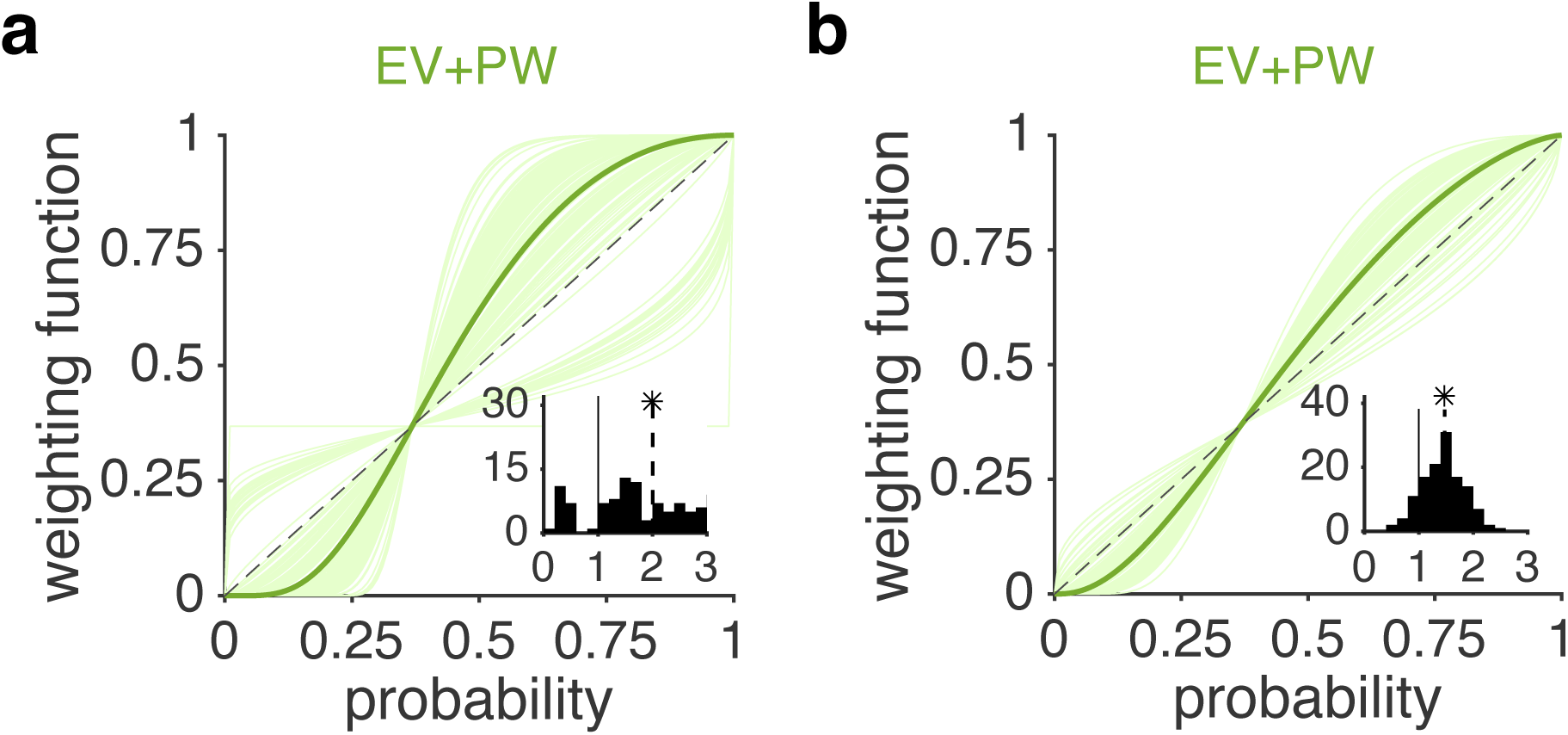
Estimated utility and probability weighting functions based on the EV+PW model. (**a**) Estimated probability weighting function based on the EV+PW model for the juice-gambling task. Each curve shows the result of the fit for one session of the experiment and the thick green curve is based on the median of the fitting parameter. The dashed line is the unity line. The inset shows the distribution of estimated parameters for the probability weighting function (*γ*). The dashed lines show the medians, and a star indicates that the median of the distribution is significantly different from 1 (two-sided sign-test; *p <* 0.05). (**b**) The same as in (a) but for the token-gambling task.

**Figure 11-1.**
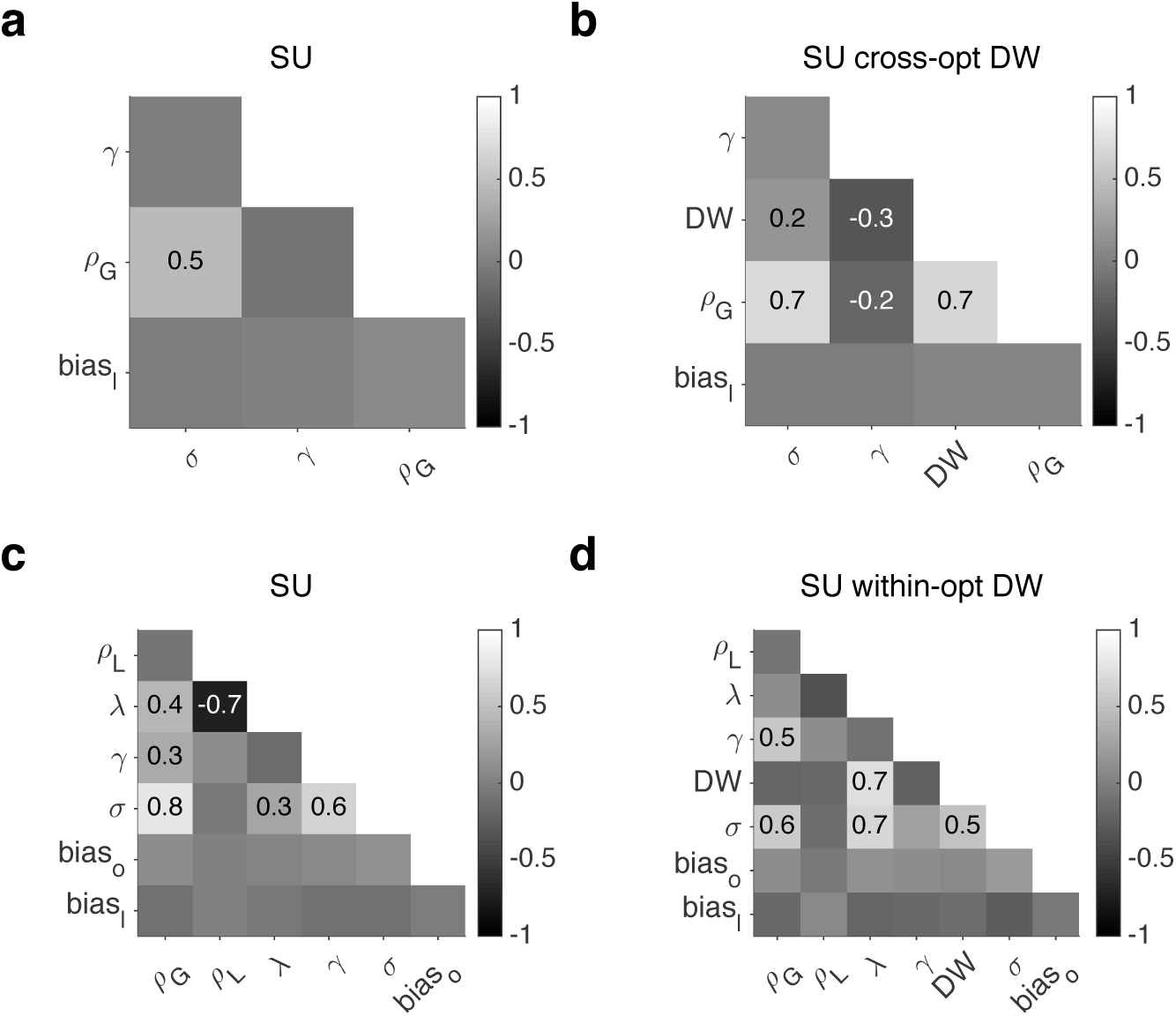
Correlation between model parameters using the inverse of the Hessian matrix. The matrix of correlation coefficients between a given model parameters is calculated from the inverse of the Hessian matrix. (**a-b**) Color of each square indicates the correlation coefficient between each pair of parameters for the SU (a) and the SU with DW models (b) during the juice-gambling task. Correlation coefficients are reported for values larger than 0.1 or smaller than −0.1 only. (**c-d**) The same as in (a-b) but for the token-gambling task.

**Figure 11-2.**
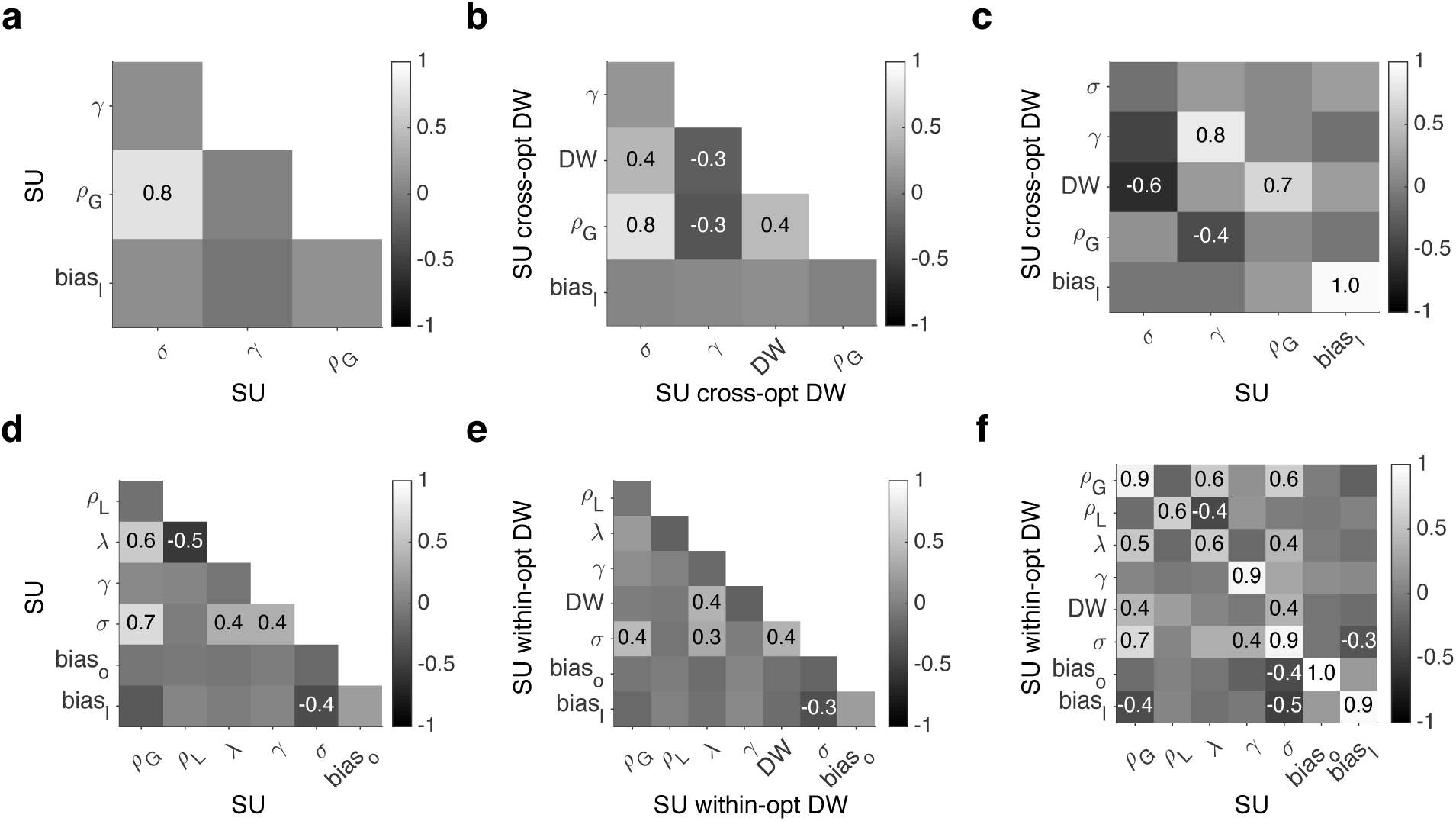
Correlation between model parameters using session-by-session estimates. For each model and between the pair of models, the matrix of correlation coefficients between model parameters is calculated using estimated model parameters in each session. (**a**-**c**) Color of each square indicates the correlation coefficient between each pair of parameters of the SU model (a), the SU with DW models (b), and between the parameters of the two models (c) during the juice-gambling task. Correlation coefficients are only reported for statistically significant values (*p* < .05, using Bonferroni correction to adjust critical p-value in each panel). (**d-f**) The same as in (a-c) but for the token-gambling task.

**Figure 11-3.**
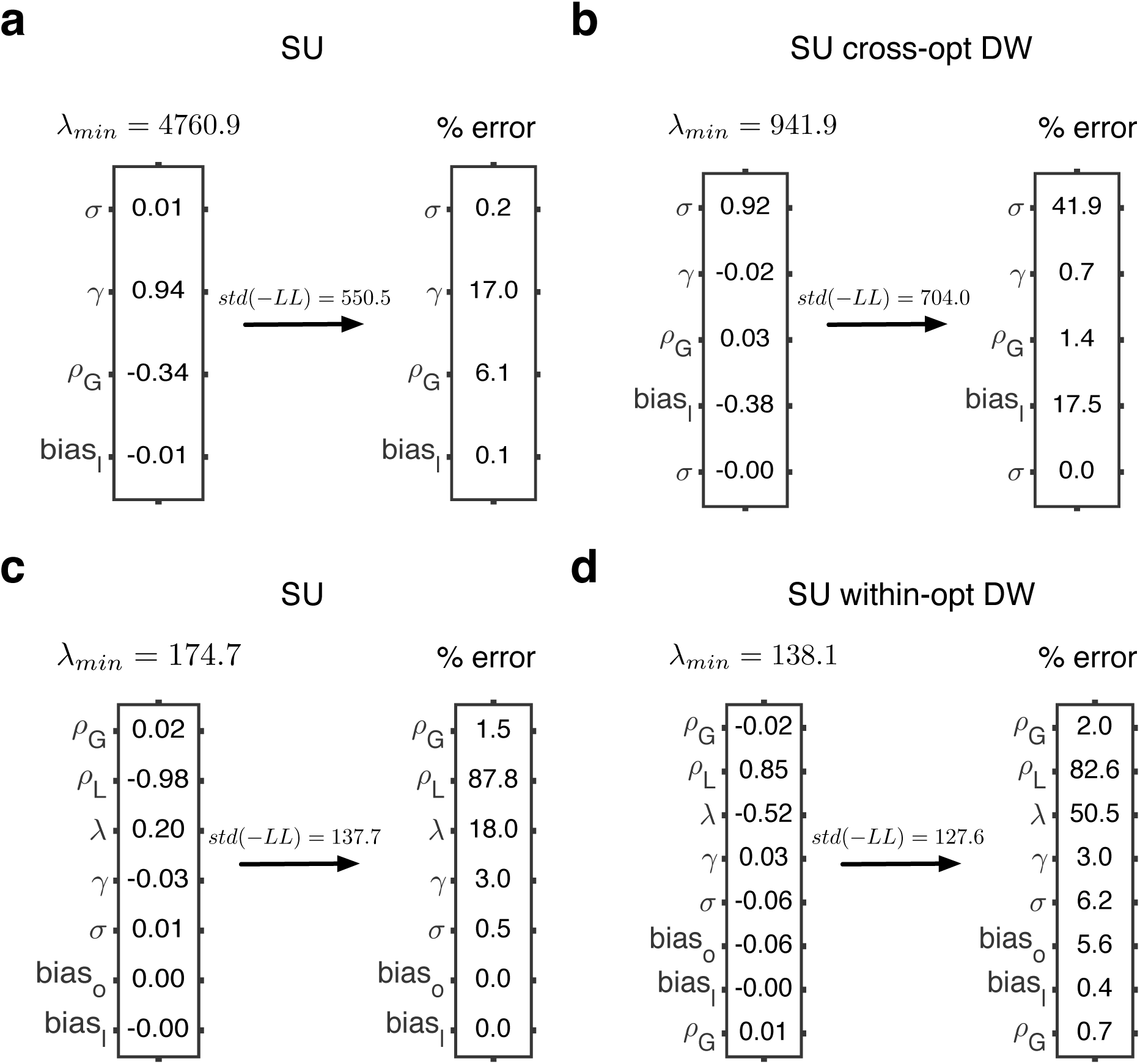
Error in the values of estimated parameters in the best models with and without differential weighting. (**a**) The left column shows the minimum eigenvalue and the corresponding eigenvector for the SU model in the juice-gambling task. The right column shows the estimated percent error for each model parameter. (**b**) The same as in (a) but for the SU model with differential weighting. (**c**-**d**) The same as in (a-b) but for the token-gambling task.

## REFERENCES

1. Abe H, Lee D (2011) Distributed Coding of Actual and Hypothetical Outcomes in the Orbital and Dorsolateral Prefrontal Cortex. Neuron 70(4):731–741.

2. Addessi E, Albano M, De Petrillo F, Laviola G, Mirolli M, Paglieri F, Adriani W (2015) Neurobiological basis of gambling: the joint contribution of psychobiology, cognitive ethology and robotics. Sistemi intelligenti 27(3):561–614.

3. Azab H, Hayden BY (2017) Correlates of decisional dynamics in the dorsal anterior cingulate cortex. PLoS Biology 15(11):e2003091.

4. Azab H, Hayden, BY (2018) Correlates of economic decisions in the dorsal and subgenual anterior cingulate cortices. Eur J Neurosci.

5. Bateson M, Kacelnik A (1996) Rate currencies and the foraging starling: the fallacy of the averages revisited. Behav Ecol 7(3):341–352.

6. Beran MJ, Perdue BM, Smith DJ (2014) What are my chances? Closing the gap in uncertainty monitoring between rhesus monkeys (Macaca mulatta) and capuchin monkeys (Cebus apella). J Exp Psychol Anim Learn Cogn 40(3):303–316.

7. Blanchard, TC, Hayden BY (2015) Monkeys are more patient in a foraging task than in a standard intertemporal choice task. PLoS One 10(2):e0117057.

8. Blanchard TC, Hayden BY, Bromberg-Martin ES (2015a) Orbitofrontal cortex uses distinct codes for different choice attributes in decisions motivated by curiosity. Neuron 85(3):602–614.

9. Blanchard TC, Strait CE, Hayden BY (2015b) Ramping ensemble activity in dorsal anterior cingulate neurons during persistent commitment to a decision. J Neurophysiol 114(4):2439–2449.

10. Blanchard TC, Wilke A, Hayden BY (2014) Hot-Hand Bias in Rhesus Monkeys. J Exp Psychol Anim Learn Cogn 40(3):280–286.

11. Blanchard TC, Wolfe LS, Vlaev I, Winston JS, Hayden BY (2014) Biases in preferences for sequences of outcomes in monkeys. Cognition 130(3):289–299.

12. Busemeyer JR, Townsend JT (1993) Decision field theory: A dynamic-cognitive approach to decision making in an uncertain environment. Psychol Rev 100(3):432–459.

13. Camilleri R, Newell BR (2013) Mind the gap? Description, experience, and the continuum of uncertainty in risky choice. Prog Brain Res 202:55–71.

14. Caraco T, Martindale S, Whittam TS (1980) An Empirical Demonstration of Risk-Sensitive Foraging Preferences. Anim Behav 28:820–830.

15. Chen MK, Lakshminarayanan V, Santos LR (2006) How basic are behavioral biases? Evidence from capuchin monkey trading behavior. J Political Econ 114(3):517–537.

16. Chen X, Stuphorn V (2015) Sequential selection of economic good and action in medial frontal cortex of macaques during value-based decisions. Elife 4:e09418.

17. De Petrillo F, Ventricelli M, Ponsi GPA, Addessi E (2015) Do tufted capuchin monkeys play the odds? Flexible risk preferences in Sapajus spp. Anim Cogn 18(1):119–130.

18. Diamond, A. (2013) Executive functions. Annu Rev Psychol 64:135–168.

19. Egan, LC, Santos, LR, Bloom P (2007) The origins of cognitive dissonance: evidence from children and monkeys. Psychol Sci 18:978–983.

20. Farashahi S, Donahue CH, Khorsand P, Seo H, Lee D, Soltani A (2017) Metaplasticity as a Neural Substrate for Adaptive Learning and Choice under Uncertainty. Neuron 94(2):401–414.

21. Hayden BY (2016) Time discounting and time preference in animals: A critical review. Psychon Bull Rev 23(1):39–53.

22. Hayden BY, Heilbronner SR, Nair AC, Platt ML (2008) Cognitive influences on risk-seeking by rhesus macaques. Judgm Decis Mak 3(5):389–395.

23. Hayden BY, Heilbronner SR, Platt ML (2010) Ambiguity aversion in rhesus macaques. Front Neurosci 4:166.

24. Hayden BY, Pearson JM, Platt ML (2009) Fictive Reward Signals in the Anterior Cingulate Cortex. Science 324:948–950.

25. Hayden BY, Platt ML (2007) Temporal Discounting Predicts Risk Sensitivity in Rhesus Macaques. Curr Biol 17(1):49–53.

26. Hayden BY, Platt ML (2009) Gambling for Gatorade: risk-sensitive decision making for fluid rewards in humans. Anim Cogn, 12(1):201–207.

27. Heilbronner SR (2017) Modeling risky decision-making in nonhuman animals: shared core features. Current opinion in behavioral sciences 16:23–29.

28. Heilbronner SR, Hayden BY (2013) Contextual factors explain risk-seeking preferences in rhesus monkeys. Front Neurosci 7:1–7

29. Heilbronner SR, Hayden BY (2016) The description-experience gap in risky choice in nonhuman primates. Psychon Bull Rev 23(2):593–600.

30. Heilbronner SR, Rosati AG, Stevens JR, Hare B, Hauser MD (2008) A fruit in the hand or two in the bush? Divergent risk preferences in chimpanzees and bonobos. Biol Lett 4(3):246–249

31. Hertwig R, Barron G, Weber EU, Erev I (2004) Decisions from Experience and the Effect of Rare Events in Risky Choice. Psychol Sci 15(8):534–539.

32. Hertwig R, Erev I (2009) The description–experience gap in risky choice. Trends Cogn Sci 13(12):517–523.

33. Kacelnik A, Bateson M (1997) Risk-sensitivity: crossroads for theories of decision-making. Trends Cogn Sci 1(8):304–309.

34. Kahneman D, Tversky A. (1979) Prospect Theory: An Analysis of Decision under Risk. Econometrica 47(2):263–292.

35. Kahneman D, Tversky A (2000) Choices, values, and frames. In Handbook of the Fundamentals of Financial Decision Making, pp269–278. Cambridge: Cambridge UP.

36. Lakshminarayanan VR, Chen MK, Santos LR (2011) The evolution of decision-making under risk: Framing effects in monkey risk preferences. J Exp Soc Psychol 47(3):689–693.

37. Lichtenstein S, Slovic P (2006) The construction of preference. Cambridge: Cambridge UP.

38. Ludvig EA, Madan CR, Pisklak JM, Spetch ML (2014) Reward context determines risky choice in pigeons and humans. Biol Lett 10(8)

39. Ludvig EA, Spetch ML (2011) Of Black Swans and Tossed Coins: Is the Description-Experience Gap in Risky Choice Limited to Rare Events? PloS One 6(6):e20262.

40. Marshall JAR, Trimmer PC, Houston AI, McNamara JM (2013) On evolutionary explanations of cognitive biases. Trend Ecol Evol 28(8):469–473.

41. McCoy AN, Platt ML (2005) Risk-sensitive neurons in macaque posterior cingulate cortex. Nat Neurosci 8(9):1220.

42. Mendelson TC, Fitzpatrick CL, Hauber ME, Pence CH, Rodríguez RL, Safran RJ, Stevens JR. (2016) Cognitive Phenotypes and the Evolution of Animal Decisions. Trend Ecol Evol 31(11):850–859.

43. O’Neill M, Schultz W (2010) Coding of reward risk by orbitofrontal neurons is mostly distinct from coding of reward value. Neuron 68(4):789–800.

44. Paglieri F, Addessi E, De Petrillo F, Laviola G, Mirolli M, Parisi D, Petrosino G, Ventricelli M, Zoratto F, Adriani W (2014) Nonhuman gamblers: lessons from rodents, primates, and robots. Front Behav Neurosci 8(33):22–60.

45. Pearson JM, Hayden BY, Platt ML (2010) Explicit information reduces discounting behavior in monkeys. Front Psychol 1:237.

46. Pearson JM, Watson KK, Platt ML (2014) Decision Making: The Neuroethological Turn. Neuron 82(5):950–965.

47. Platt ML, Huettel SA (2008) Risky business: the neuroeconomics of decision making under uncertainty. Nat Neurosci 11(4):398–403.

48. Roe RM, Busemeyer JR, Townsend JT (2001) Multialternative decision field theory: A dynamic connectionist model of decision making. Psychol Rev 108(2):370–392.

49. Santos LR, Rosati AG (2015) The Evolutionary Roots of Human Decision Making. Annu Rev of Psychol 66:321–347.

50. Shimojo S, Simion C, Shimojo E, Scheier C (2003) Gaze bias both reflects and influences preference. Nat Neurosci 6(12):1317–1322.

51. Seo H, and Lee D (2009) Behavioral and neural changes after gains and losses of conditioned reinforcers. J Neurosci 29(11):3627–3641.

52. Seo H, Cai X, Donahue CH, Lee D (2014). Neural correlates of strategic reasoning during competitive games. Science Sep 18:1256254.

53. So NY, Stuphorn V (2010) Supplementary eye field encodes option and action value for saccades with variable reward. J Neurophysiol 104(5):2634–2653.

54. So NY, Stuphorn V (2012) Supplementary eye field encodes reward prediction error. J Neurosci 32(9):2950–2963.

55. Stauffer WR, Lak A, Bossaerts P, Schultz W (2015) Economic Choices Reveal Probability Distortion in Macaque Monkeys. J Neurosci 35(7):3146–3154.

56. Stephens DW, Anderson D (2001) The adaptive value of preference for immediacy: when shortsighted rules have farsighted consequences. Behav Ecol 12(3):330–339.

57. Stevens JR, Rosati AG, Ross KR, Hauser MD (2005) Will Travel for Food: Spatial Discounting in Two New World Monkeys. Curr Biol 15(20):1855–1860.

58. Strait CE, Blanchard, TC, Hayden BY (2014) Reward Value Comparison via Mutual Inhibition in Ventromedial Prefrontal Cortex. Neuron 82(6):1357–1366.

59. Strait CE, Sleezer, BJ, Blanchard, TC, Azab H, Castagno MD, Hayden BY (2016) Neuronal selectivity for spatial positions of offers and choices in five reward regions. J Neurophysiol 115(3):1098–1111.

60. Strait CE, Sleezer BJ, Hayden BY (2015) Signatures of Value Comparison in Ventral Striatum Neurons. PLoS Biol 13(6)

61. Strier KB (2016) Primate behavioral ecology. New York: Routledge.

62. Trepel C, Fox CR, Poldrack RA (2005) Prospect theory on the brain? Toward a cognitive neuroscience of decision under risk. Cogn Brain Res 23:34–50.

63. Yamada H, Tymula A, Louie K, Glimcher PW (2013) Thirst-dependent risk preferences in monkeys identify a primitive form of wealth. Proc Natl Acad Sci 110(39):15788–15793.

